# Computational noise in reward-guided learning drives behavioral variability in volatile environments

**DOI:** 10.1101/439885

**Authors:** Charles Findling, Vasilisa Skvortsova, Rémi Dromnelle, Stefano Palminteri, Valentin Wyart

## Abstract

When learning the value of actions in volatile environments, humans often make seemingly irrational decisions which fail to maximize expected value. We reasoned that these ‘non-greedy’ decisions, instead of reflecting information seeking during choice, may be caused by computational noise in the learning of action values. Here, using reinforcement learning (RL) models of behavior and multimodal neurophysiological data, we show that the majority of non-greedy decisions stems from this learning noise. The trial-to-trial variability of sequential learning steps and their impact on behavior could be predicted both by BOLD responses to obtained rewards in the dorsal anterior cingulate cortex (dACC) and by phasic pupillary dilation – suggestive of neuromodulatory fluctuations driven by the locus coeruleus-norepinephrine (LC-NE) system. Together, these findings indicate that most of behavioral variability, rather than reflecting human exploration, is due to the limited computational precision of reward-guided learning.

## Introduction

In uncertain environments, decision-makers learn rewarding actions by trial-and-error to maximize their expected payoff (Fig. 1a). An important challenge is that reward contingencies typically change over time, and thus a less-rewarded action at a given point in time can become more rewarding later (Fig. 1b). Versatile machine learning algorithms, known collectively as ‘reinforcement learning’ (RL), describe the changing values of possible actions and the policy used to choose among them^1^. One biologically plausible class of RL models updates the expected values associated with possible actions sequentially based on the ‘prediction error’ (PE) between obtained and expected reward – a learning scheme known as the Rescorla-Wagner rule^2^. At any given time point, the decision-maker chooses based on the difference in expected value between possible actions, by selecting the action associated with the largest expected reward. However, in volatile environments in which reward contingencies change rapidly over time, human decision-makers make a substantial number of seemingly irrational decisions which do not maximize the expected value predicted by reinforcement learning^3,4^. These decisions are often coined as ‘non-greedy’ in contrast to value-maximizing, ‘greedy’ decisions.

**Figure 1.**
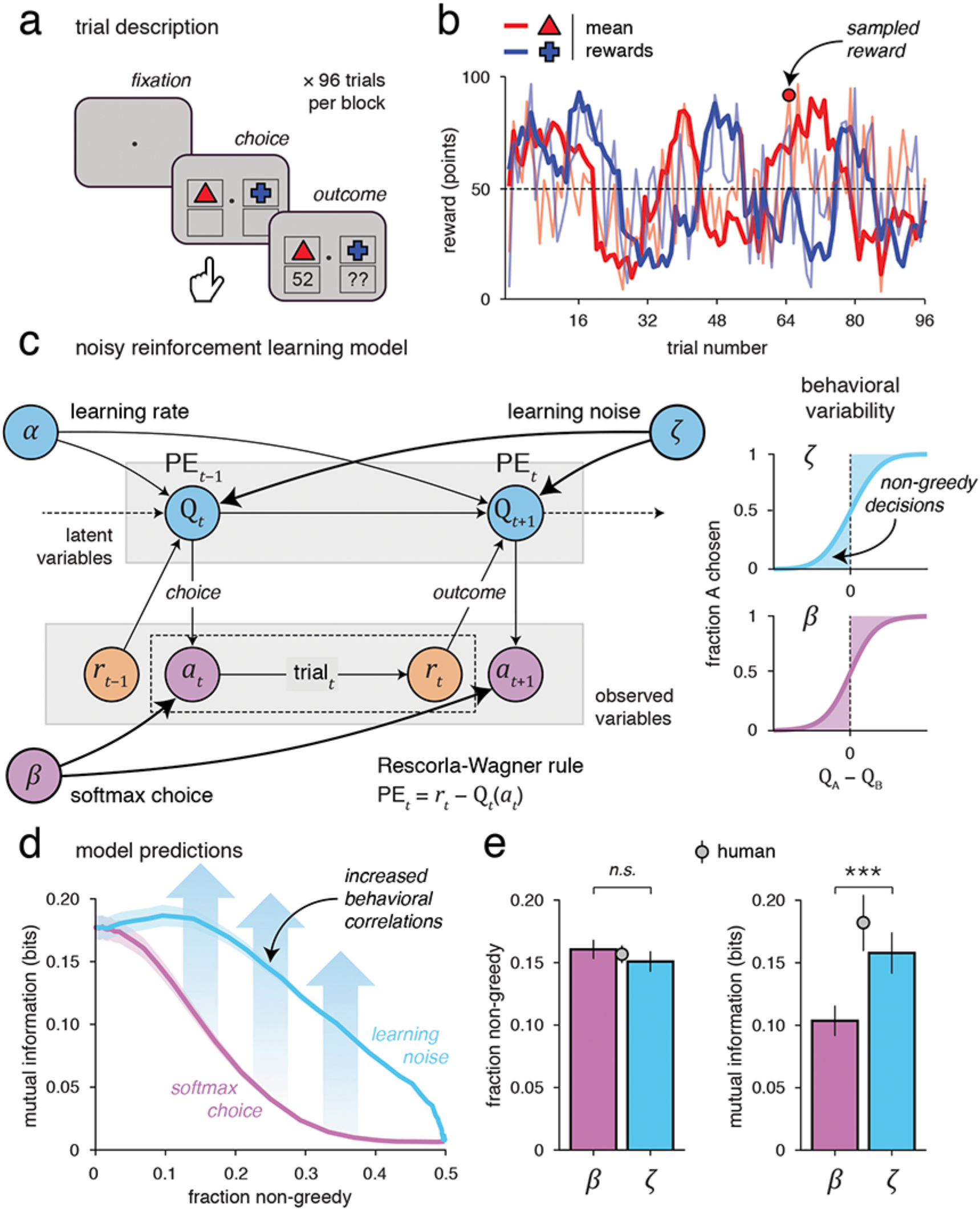
Experimental paradigm and noisy reinforcement learning model. (**a**) Trial structure in the restless, two-armed bandit task divided into short blocks of trials. On every trial, participants were asked to choose one of two reward sources depicted by colored shapes, and then observed its associated outcome (from 1 to 99 points, converted into real financial incentives at the end of the experiment). (**b**) Example of drifts in the magnitude of rewards that can be obtained from the two sources. Rewards were sampled from probability distributions whose means drifted independently across trials. Thick lines represent the drifting means of the two probability distributions, whereas thin lines correspond to reward samples drawn from the probability distributions that can be obtained if chosen on each trial. (**c**) Graphical representation of the noisy RL model used to fit human behavior in the task. The Rescorla-Wagner learning rule applied to update action values is corrupted by additive random noise. The choice process is modeled using a stochastic ‘softmax’ action selection policy. Learning noise is assumed to be negligible in exact RL models. Inset: illustration of the fraction of non-greedy decisions predicted either by an exact RL model followed by a softmax action selection policy (top panel, area shaded in purple), or by a noisy RL model followed by a purely value-maximizing action selection policy (bottom panel, area shaded in blue). Exact and noisy RL models are indistinguishable based only on their predicted fraction of non-greedy decisions. (**d**) Predicted relationship between the fraction of non-greedy decisions and the mutual information of successive decisions for the exact (purple) and noisy (blue) RL models. For the same fraction of non-greedy decisions, a noisy RL model predicts larger behavioral correlations (mutual information) across successive decisions than an exact RL model. (**e**) Falsification of the exact RL model through model simulations. Simulated (bars) and observed (dot) fraction of non-greedy decisions (left panel) and mutual information of successive decisions (right panel). While the overall fraction of non-greedy decisions is well captured by both noisy and exact RL models, the observed mutual information is predicted more accurately by simulations of noisy RL than exact RL. Error bars correspond to s.e.m.

A prominent hypothesis regarding the source of these non-greedy decisions is that they are the result of a compromise during choice between exploiting a currently well-valued action vs. exploring other, possibly better-valued actions – known as the ‘exploration-exploitation’ trade-off. In this view, information seeking motivates non-greedy decisions. Indeed, for a value-maximizing agent, lower-valued actions are selected less often and thus their expected values are more uncertain than those of higher-valued actions. Non-greedy decisions in favor of recently unchosen actions thus effectively reduce uncertainty about their current value and increase long-term payoff^4–7^. Different regions of the human frontal cortex have been shown to activate during non-greedy decisions in volatile environments, including the dorsal anterior cingulate cortex (dACC) and the frontopolar cortex (FPC)^3,8^. In agreement with an ‘information seeking’ account of non-greedy decisions, these two regions have been linked with uncertainty monitoring. Specifically, dACC activity correlates with the volatility of reward contingencies and increases during uncertain decisions between similarly-valued actions^9–11^, whereas FPC activity reflects the expected value of unchosen actions^12–14^. An important corollary of this view is that non-greedy decisions are driven solely by adaptive regulations of the choice process: both stochastic regulations, such as the use of a ‘soft’ value-maximizing policy, and deterministic regulations, including ‘information bonuses’ added to the value of unchosen actions^7,15^. By contrast, the reinforcement learning process which tracks action values over time is implicitly assumed to follow exactly the hypothesized Rescorla-Wagner learning rule after each obtained reward.

However, it has recently been shown that the accuracy of human perceptual decisions based on multiple sensory cues is bounded not by variability in the choice process, but rather by ‘inference’ noise arising during evidence accumulation^16,17^. This computational noise is responsible for a dominant fraction of non-greedy decisions which do not follow the optimal Bayes rule of probabilistic reasoning. An intriguing possibility is that the learning process at the heart of reward-guided decision-making might be subject to the same kind of computational noise – in this case, random variability in the update of action values (Fig. 1c). Critically, the existence of intrinsic noise in reinforcement learning would trigger non-greedy decisions due to random deviations between exact applications of the learning rule and its noisy realizations following each obtained reward. This noise-driven source of non-greedy decisions is not mutually exclusive with information seeking: non-greedy decisions can be driven simultaneously by learning noise and information seeking. In other words, in this view, an unknown fraction of non-greedy decisions results not from overt information seeking during choice, as assumed by existing theories and computational models, but from the limited precision of the underlying reinforcement learning process which is classically assumed to be infinite.

To determine whether, and to what extent, learning noise drives non-greedy decisions during reward-guided decision-making, we first derived a theoretical formulation of RL which allows for random noise in its core computations. In a series of behavioral and neuroimaging experiments, tested over a total of 90 human participants, we then quantified the fraction of non-greedy decisions which could be attributed to learning noise, and identified its neurophysiological substrates using functional magnetic resonance imaging (fMRI) and pupillometric recordings.

## Results

### Experimental protocol and computational model

We designed a canonical restless, two-armed bandit game. Over three experiments, a total of 90 human participants were asked to maximize their monetary payoff by sampling repeatedly from one among two reward sources depicted by colored shapes (see Methods). Experiments were divided into short blocks of trials, each involving a new pair of colored shapes. On each trial, participants were asked to choose one of the two shapes, and then observed its associated outcome (Fig. 1a). The payoffs that could be obtained from either shape (from 1 to 99 points, converted into real financial incentives at the end of the experiment) were sampled from probability distributions whose means drifted independently across trials – thereby encouraging participants to track these mean values over the course of each block (Fig. 1b).

To characterize the origin of non-greedy decisions made in this task, we derived a RL model in which the Rescorla-Wagner rule applied to update action values Q_*t*_ is corrupted by additive random noise *ε*_*t*_ (Fig. 1c):

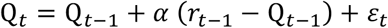

where *α* is the learning rate used to update action values based on the prediction error (PE) between obtained reward *r*_*t*-1_ and expected reward Q_*t*-1_ on the previous trial, and *ε*_*t*_ is drawn from a normal distribution with zero mean and standard deviation *σ*_*t*_ equal to a constant fraction *ζ* of the magnitude of the PE:

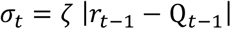

This ‘multiplicative’ structure of the noise, which assumes larger deviations during larger updates of action values, follows the ubiquitous Weber’s law of intensity sensation prevalent in numerous perceptual domains (including vision, numerosity and time), and in the magnitude of associated neural responses^18–20^.

As in existing theories, we modeled the choice process using a stochastic ‘softmax’ action selection policy, controlled by an ‘inverse temperature’ *β*. Importantly, although learning noise and choice stochasticity both generate non-greedy decisions as defined by exact (noise-free) RL (Fig. 1c), the two sources of behavioral variability make different predictions regarding the temporal structure of decisions across successive trials. Indeed, learning noise corrupts the action values which are gradually updated across trials, and used to drive successive decisions. By contrast, choice stochasticity reads out action values without altering them and is independently distributed across trials. Therefore, for the same fraction of non-greedy decisions simulated either using learning noise or choice stochasticity, learning noise engenders larger behavioral correlations across successive decisions (Fig. 1d).

### Dominant contribution of learning noise to non-greedy decisions

Looking at the first, neuroimaging experiment (experiment 1, *N* = 29), participants selected the shape associated with the largest mean payoff on a majority of trials (64.9 ± 0.9%, mean ± s.e.m., *t*-test against chance: *t*_28_ = 17.2, *p* < 0.001). As anticipated, participants also made a substantial fraction of non-greedy decisions – which do not maximize expected value with respect to an exact (noise-free) RL model (15.7 ± 0.7%).

We next performed Bayesian Model Selection (BMS) to quantify the contributions of learning- and choice-driven sources of variability to non-greedy decisions (Fig. 2a). Using particle filtering procedures to obtain estimates of model evidence conditioned on human decisions (see Methods), we found that a RL model corrupted by learning noise explained human behavior significantly better than an exact RL model (*ζ* fitted vs. *ζ* = 0, fixed-effects: BF ≈ 10^12.9^, random-effects: exceedance *p* = 0.941). This first finding indicates that the sequential updating of action values is subject to a significant amount of noise. A softmax action selection policy also outperformed a purely value-maximizing, ‘argmax’ policy (*β* fitted vs. *β* → ∞, fixed-effects: BF ≈ 10^44.5^, random-effects: exceedance *p* > 0.999) – thereby indicating that non-greedy decisions are driven both by learning noise and choice stochasticity (Supplementary Fig. 1a). This behavioral pattern was fully replicated in the second experiment (experiment 2, *N* = 30): like participants tested in the first experiment, participants featured both learning noise (fixed-effects: BF ≈ 10^37.3^, random-effects: exceedance *p* > 0.999) and a softmax action selection policy (fixed-effects: BF ≈ 10^37.9^, random-effects: exceedance *p* > 0.999; Supplementary Fig. 1b). To validate these findings, we implemented a ‘model recovery’ procedure, which confirmed that our BMS procedure was capable of correctly distinguishing learning noise from choice stochasticity in our task (Fig. 2b).

**Figure 2.**
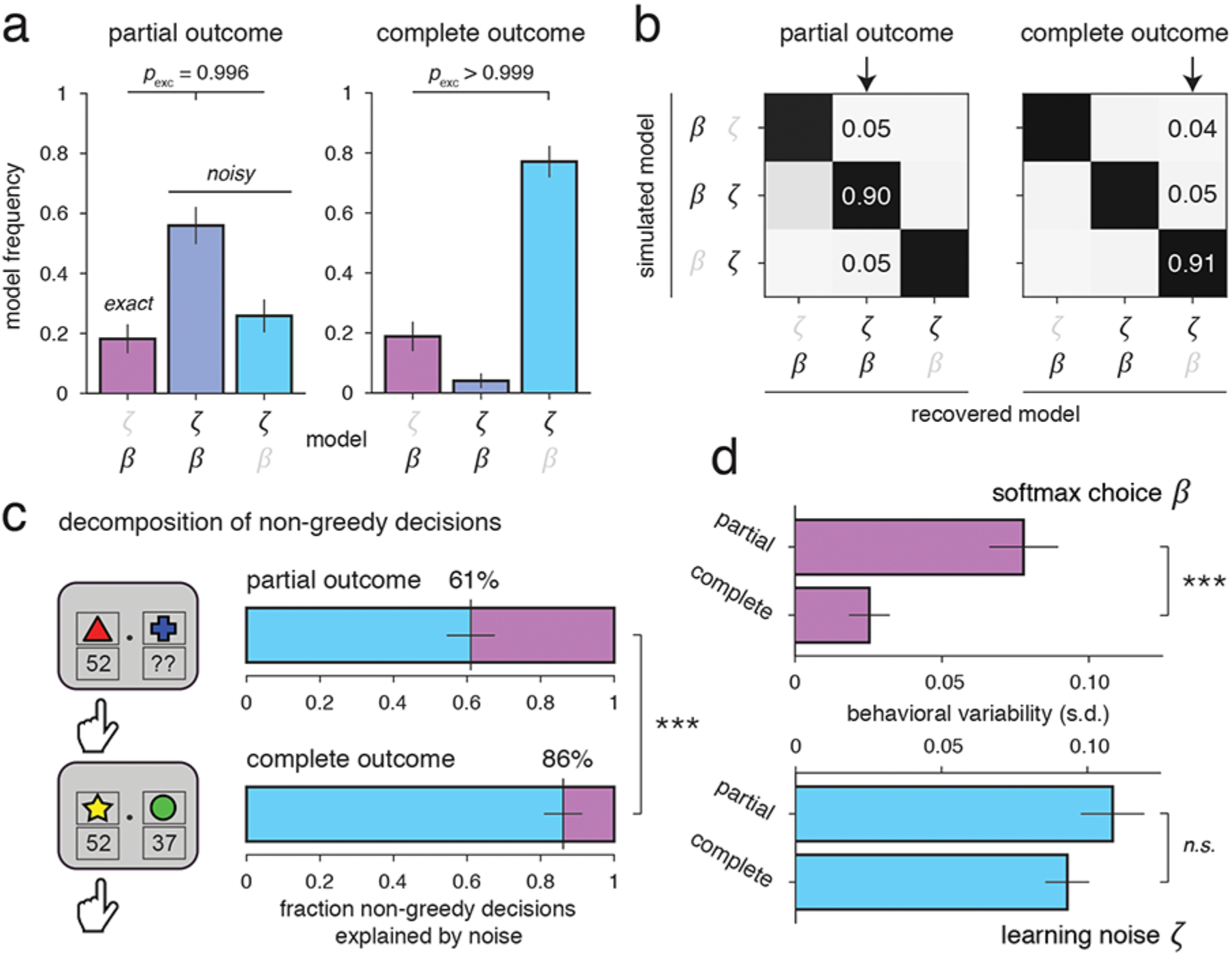
Contributions of learning noise and choice stochasticity to non-greedy decisions. (**a**) Bayesian model selection results in the partial outcome condition (left panel) and the complete outcome condition (right panel). Estimated model frequencies for exact RL (β, left bar), noisy RL with a softmax action selection rule (β and ζ, middle bar) and noisy RL with an argmax action selection rule (ζ, right bar). Noisy RL outperforms exact RL in both outcome conditions. Error bars correspond to s.d. of the estimated Dirichlet distribution. (**b**) Model recovery results in the partial outcome condition (left panel) and the complete outcome condition (right panel). Confusion matrices displaying the estimated model frequencies of exact and noisy RL (columns) obtained for simulations of exact and noisy RL (rows). Bayesian model selection is able to recover accurately the simulated RL model in both outcome conditions. (**c**) Fraction of non-greedy decisions explained by learning noise (blue area) and choice stochasticity (purple area) in the partial outcome condition (top panel) and the complete outcome condition (bottom panel). Learning noise alone explains more than half of non-greedy decisions in the partial outcome condition, and almost all of non-greedy decisions in the complete outcome condition. (**d**) Amounts of behavioral variability (expressed as standard deviation on the difference between action values predicted by the noisy RL model) due separately to learning noise (bottom panel) and choice stochasticity (top panel) in the partial and complete outcome conditions. Noise-driven variability is not different between the two outcome conditions, whereas choice-driven variability is strongly reduced in the complete outcome condition. Error bars correspond to s.e.m. Three stars correspond to a significant difference at p < 0.001, n.s. to a non-significant difference.

Importantly, the exact RL model could be falsified by comparing the sequential dependency of its simulated decisions to the sequential dependency of human decisions (see Methods). While the overall fraction of non-greedy decisions was well captured by both noisy and exact RL models (human: 15.7 ± 0.7%; noisy RL: 15.0 ± 0.8%; exact RL: 16.0 ± 0.7%), the mutual information of successive human decisions was better predicted by simulations of noisy RL than exact RL (human: 0.126 ± 0.016 bit; noisy RL: 0.111 ± 0.011 bit; exact RL: 0.073 ± 0.008 bit; paired *t*-test, *t*_28_ = 5.8, *p* < 0.001; Fig. 1e). This behavioral signature allows us to falsify choice stochasticity as the dominant source of non-greedy decisions in this canonical task^21^.

Given the presence of both sources of behavioral variability, we went further and quantified the respective contributions of learning noise and choice stochasticity to non-greedy decisions. For this purpose, we first estimated the trial-to-trial trajectories of latent action values corrupted by learning noise conditioned on observed human decisions in every block (see Methods). We then assessed the fraction of non-greedy decisions that could be uniquely attributed to learning noise – i.e., trials in which noisy realizations of the learning rule resulted in an opposite ranking of action values to exact applications of the same rule. This quantitative analysis revealed that learning noise alone explained as much as 60.6 ± 6.6% of non-greedy decisions (Fig. 2c). Again, we replicated this pattern in the second experiment (65.6 ± 6.0%). In contrast to existing accounts, this pattern of findings indicates that behavioral variability is driven to a large part by random noise in the update of action values, rather than by stochasticity in the choice process.

### Dissociating learning noise from information seeking

We then sought to dissociate the observed learning noise from information seeking. One obvious way consists in showing that the behavioral variability stemming from learning noise is not aimed explicitly at seeking information about recently unchosen actions – whose associated rewards have not been observed and are thus uncertain. To test this important prediction, we contrasted in both experiments the classical ‘partial outcome’ condition in which participants observe only the reward yielded by the selected shape (Fig. 2c, top panel), with another ‘complete outcome’ condition in which participants additionally observe the foregone reward which would have been obtained if the other, unchosen shape had been selected^12,22^ (Fig. 2c, bottom panel). In this additional condition, performed by the same participants, there is by definition no incentive to explore – i.e., to choose actions which do not maximize expected value – given that there is equal uncertainty about the values of chosen and unchosen actions. Therefore, the residual behavioral variability observed in the complete outcome condition, if any, should be entirely driven implicitly by learning noise and not explicitly by a softmax action selection policy.

Before testing this prediction, we first verified that participants used the information about foregone actions to select the shape associated with the largest mean payoff more often than in the partial outcome condition (experiment 1, partial: 64.9 ± 0.9%, complete: 70.3 ± 0.7%, paired *t*-test, *t*_28_ = 4.9, *p* < 0.001; experiment 2, partial: 64.9 ± 0.8%, complete: 69.5 ± 1.1%, paired *t*-test, *t*_29_ = 3.5, *p* = 0.002). Model fitting confirmed that participants used the reward from the unchosen action to update its associated value (fixed-effects: BF ≈ 10^11.7^, random-effects: exceedance *p* > 0.999). Interestingly, the learning rate *α* associated with the unchosen action was not different from the one associated with the chosen action (chosen: 0.596 ± 0.043, unchosen: 0.621 ± 0.042, paired *t*-test, *t*_28_ = 1.5, *p* = 0.156, BF_H0_ = 2.0), consistent with the idea that participants learnt equally from obtained and foregone rewards^22–24^.

Interestingly, participants made a lower but still substantial amount of non-greedy decisions in the complete outcome condition (experiment 1, partial: 15.7 ± 0.7%, complete: 11.9 ± 0.7%, paired *t*-test, *t*_28_ = −4.2, *p* < 0.001; experiment 2, partial: 16.5 ± 1.0%, complete: 12.9 ± 0.9%, paired t-test, *t*_29_ = −3.1, *p* = 0.004) – consistent with the presence of learning noise in this condition. BMS confirmed this prediction by showing that a noisy RL model explained human behavior decisively better than an exact RL model in the complete outcome condition (fixed-effects: BF ≈ 10^40.2^, randomeffects: exceedance *p* > 0.999). Furthermore, in contrast to what was observed in the partial outcome condition, BMS indicated that a purely value-maximizing argmax action selection policy fitted the choice process decisively better than an exploratory softmax policy (fixed-effects: BF ≈ 10^10.6^, random-effects: exceedance *p* > 0.999, Supplementary Fig. 2a). Consequently, the split of non-greedy decisions in this condition showed that learning noise explained almost all of non-greedy decisions (86.1 ± 5.2%, complete vs. partial: paired *t*-test, *t*_28_ = 3.6, *p* = 0.001, Fig. 2c, bottom panel), a pattern fully replicated in the second experiment (89.1 ± 3.9%, complete vs. partial: paired *t*-test, *t*_29_ = 3.3, *p* = 0.003).

We further confirmed that this increased fraction of non-greedy decisions explained by learning noise was due to a change in action selection policy rather than an increased learning noise. Instead of computing the relative fraction of non-greedy decisions driven by learning noise, we estimated the raw amount of behavioral variability due separately to learning noise and to choice stochasticity (Fig. 2d, see Methods). This analysis confirmed both our predictions: choice-driven variability was reduced substantially in the complete outcome condition (partial: 0.078 ± 0.010, complete: 0.025 ± 0.007, paired *t*-test, *t*_28_ = −4.6, *p* < 0.001), whereas noise-driven variability was not different across the two conditions (partial: 0.110 ± 0.010, complete: 0.093 ± 0.007, paired *t*-test, *t*_28_ = −1.2, *p* = 0.240, BF_H0_ = 2.6). Together, these findings indicate that learning noise does not aim explicitly at seeking information about recently unchosen actions, but rather reflects computational constraints on the underlying learning process.

### Dissociating learning noise from model misspecification

One important possible confound is that part of the observed learning noise would be caused not by *random* deviations around the hypothesized Rescorla-Wagner rule, but by *systematic* deviations from this canonical learning rule – in other words, a misspecification of our learning model. In particular, different participants might be using different learning rules, which would then be captured as learning noise by our noisy RL model that assumes that the Rescorla-Wagner rule is being used. To decompose learning noise into systematic and random deviations from the Rescorla-Wagner rule, we ran a third, behavioral experiment (experiment 3, *N* = 30) where we estimated the consistency of human decisions across repetitions of the same sequence of rewards. Unbeknownst to participants, we made each of them play the exact same blocks of trials twice in the complete outcome condition (Fig. 3a, see Methods).

**Figure 3.**
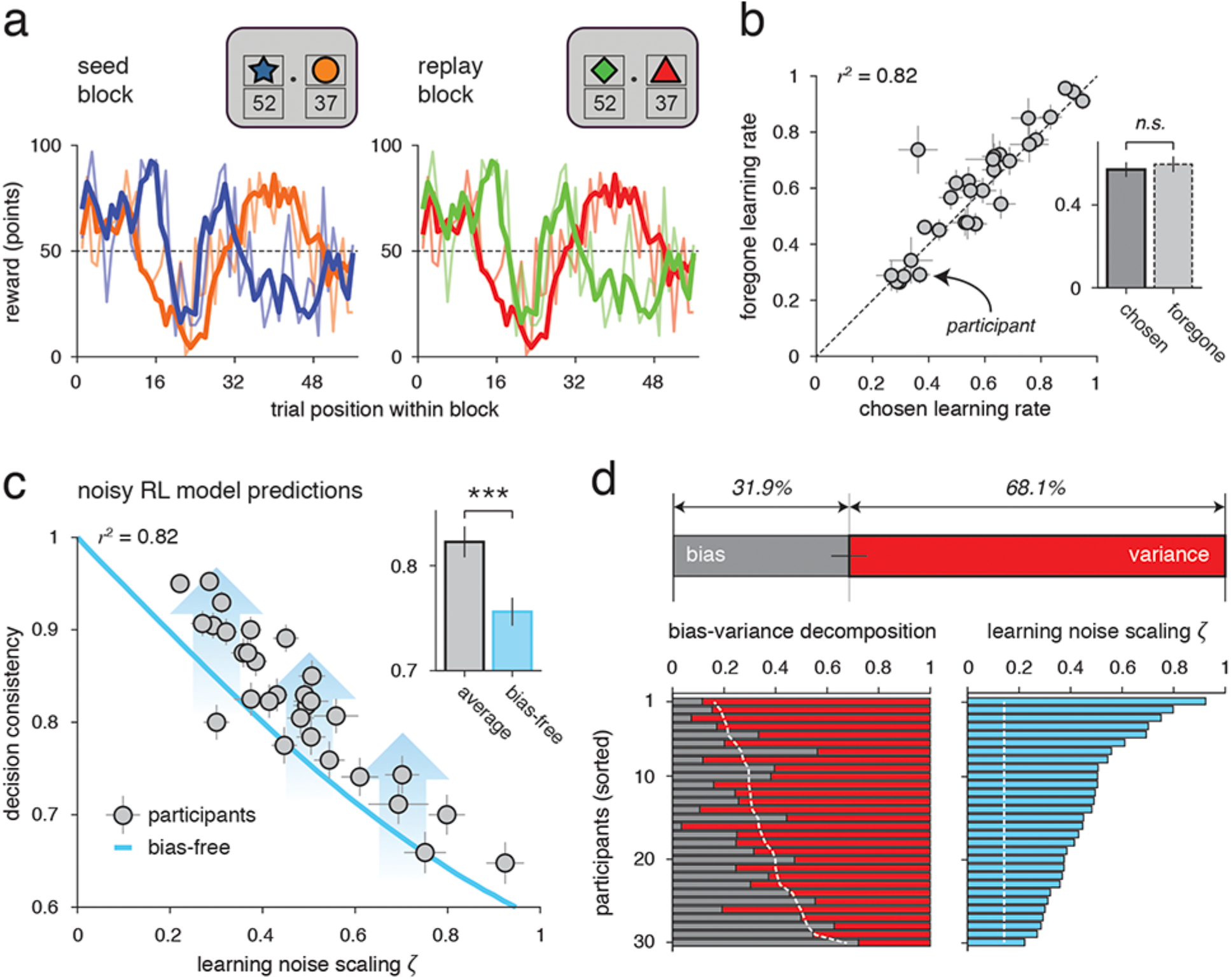
Decomposition of learning noise into ultimately predictable and unpredictable terms. (**a**) Trial structure in the experiment consisting of repeated blocks of the same sequence of rewards, tested only in the complete outcome condition. Unbeknownst to participants, each ‘seed’ block of trials (left panel) was replayed at another point in the experiment (right panel) using different colored shapes. Same conventions as in Fig. 1. (**b**) Learning rates associated with chosen and unchosen actions (dots ± error bars correspond to means ± s.d.). Learning rates did not differ between chosen and unchosen actions, indicating that action values at a given trial do not depend on choices made at earlier trials. (**c**) Decision consistency (y-axis) as a function of the scaling ζ of learning noise with the magnitude of learning steps (x-axis) fitted by the noisy RL model to each participant. The blue line corresponds to the predicted relationship between the two quantities through simulations of the noisy RL model. Variability across tested participants (dots ± error bars correspond to means ± s.d.) shows the negative correlation predicted by simulations of the noisy RL model. (**d**) Bias-variance decomposition of learning noise. Top panel: fraction of learning noise explained by predictable biases (gray area) and unpredictable variance (red area). More than two thirds of learning noise are not attributable to systematic deviations from the Rescorla-Wagner rule used in our noisy RL model. Bottom panel: relationship between the Weber scaling ζ of learning noise (right) and the bias-variance decomposition of learning noise (left) across tested participants. Sorting participants by ζ reveals a clear increase in the variance term with ζ, indicating that ζ indexes random variability in learning rather than systematic deviations from the Rescorla-Wagner rule. Error bars correspond to s.e.m. (unless indicated otherwise). Three stars correspond to a significant difference at p < 0.001, n.s. to a non-significant difference.

We restricted this experiment to the complete outcome condition for two important reasons. First, we wanted behavioral variability to be driven solely by learning noise and not by a softmax action selection policy – a pattern that we replicated in this additional dataset (noisy vs. exact RL, fixed-effects: BF ≈ 10^411.8^, random-effects: exceedance *p* > 0.999; argmax vs. softmax policy, fixed-effects: BF ≈ 10^6.3^, random-effects: exceedance *p* = 0.997). Second, we wanted the action values predicted by an exact RL model to be identical across the two repetitions of the same block, irrespectively of the decisions made in the two repeated blocks. This was ensured by the observation that, as in previous experiments, participants learnt equally from obtained (chosen) and foregone (unchosen) rewards in this additional dataset (Fig. 3b; learning rate *α*, chosen: 0.571 ± 0.036, unchosen: 0.597 ± 0.038, paired *t*-test, *t*_29_ = 1.6, *p* = 0.123, BF_H0_ = 1.7). Beyond this absence of difference in mean learning rates, validated by BMS (fixed-effects: BF ≈ 10^16.0^, random-effects: exceedance *p* > 0.999), learning rates associated with chosen and unchosen actions were also highly correlated across participants (linear correlation, *r* squared = 0.821, d.f. = 28, *p* < 0.001). This pattern of findings indicates that the action values predicted by exact RL at a given trial do not depend on decisions made at earlier trials (which are likely to differ to some extent across the two repetitions of the same block).

We then applied a recently developed statistical approach^17,21^ (see Methods) to split the overall learning noise into a predictable ‘bias’ term – reflecting systematic deviations from the Rescorla-Wagner rule, and an unpredictable ‘variance’ term – reflecting random deviations around this canonical rule. In practice, we used the consistency of decisions across repeated blocks – which ranged from 64.8% to 95.2% across participants (82.3 ± 1.5%, mean ± s.e.m.), to decompose learning noise into bias and variance terms. Indeed, systematic deviations tend to increase the consistency of decisions across repeated blocks, whereas random deviations tend to decrease the same metric. We first observed that participants with larger learning noise (i.e., a steeper scaling *ζ* of noise with prediction errors) showed lower decision consistency across repeated blocks (Fig. 3c, linear correlation, *r* squared = 0.829, d.f. = 28, *p* < 0.001). This provides direct support for our hypothesis that most of the learning noise captured by the model is due to random variability rather than to systematic deviations from the learning rule assumed by the model. Importantly, variations of learning rate across participants did not show any relationship with decision consistency (linear correlation, *r* squared = 0.008, d.f. = 28, *p* = 0.645).

We next fitted the noisy RL model to each participant, and then simulated versions of the model in which learning noise was split in two additive terms: a predictable bias term whose realizations were duplicated in the two repetitions of each block, and an unpredictable variance term whose realizations were sampled independently across repeated blocks. We varied this bias-variance trade-off from zero (fully predictable) to one (fully unpredictable) for the simulations of each participant, and found that the split that best accounted for the consistency of human decisions across repeated blocks was of 31.8 ± 3.2% for the predictable term, and 68.2 ± 3.2% for the unpredictable term (Fig. 3d). This result indicates that more than two thirds of learning noise are not attributable to misspecifications of our learning model, and supports our hypothesis that learning noise primarily reflects the limited computational precision of reward-guided learning.

### Explaining decision effects as consequences of learning noise

During sequential learning in volatile environments, humans exhibit non-greedy decisions toward recently unchosen actions, but also a tendency to repeat their previous decision over and above the value difference between possible actions. This decision effect has been described in computational terms by an additional bias in the choice process, often coined as ‘choice hysteresis’, which could have beneficial (stabilizing) properties for the decision-maker^25–30^. We realized that this effect falls naturally out of the statistical properties of learning noise, without further assumptions, when fitted using an exact (noise-free) RL model. This is due to the corruption of action values which propagates across update steps, thereby creating intrinsic temporal correlations in the noisy action values used by the decision-maker. Using an exact RL model to fit human decisions, we observed a positive choice hysteresis in both partial and complete outcome conditions (experiment 1, *t*-test against zero, partial: *t*_28_ = 2.6, *p* = 0.013; complete: *t*_28_ = 5.0, *p* < 0.001). We then verified that, as predicted, simulations of the noisy RL model fitted using an exact RL model exhibited an apparent choice hysteresis, in both conditions (*t*-test against zero, partial: *t*_28_ = 4.2, *p* < 0.001; complete: *t*_28_ = 5.6, *p* < 0.001). Critically, the choice hysteresis measured in participants correlated with the apparent choice hysteresis predicted by the noisy RL model (Fig. 4a, linear correlation, partial: *r* squared = 0.553, d.f. = 27, *p* < 0.001; complete: *r* squared = 0.425, d.f. = 27, *p* < 0.001). This finding, fully replicated in experiment 2, supports our hypothesis that choice hysteresis is not caused by an explicit bias in the choice process, but rather by learning noise which propagates through corrupted action values across successive decisions.

**Figure 4.**
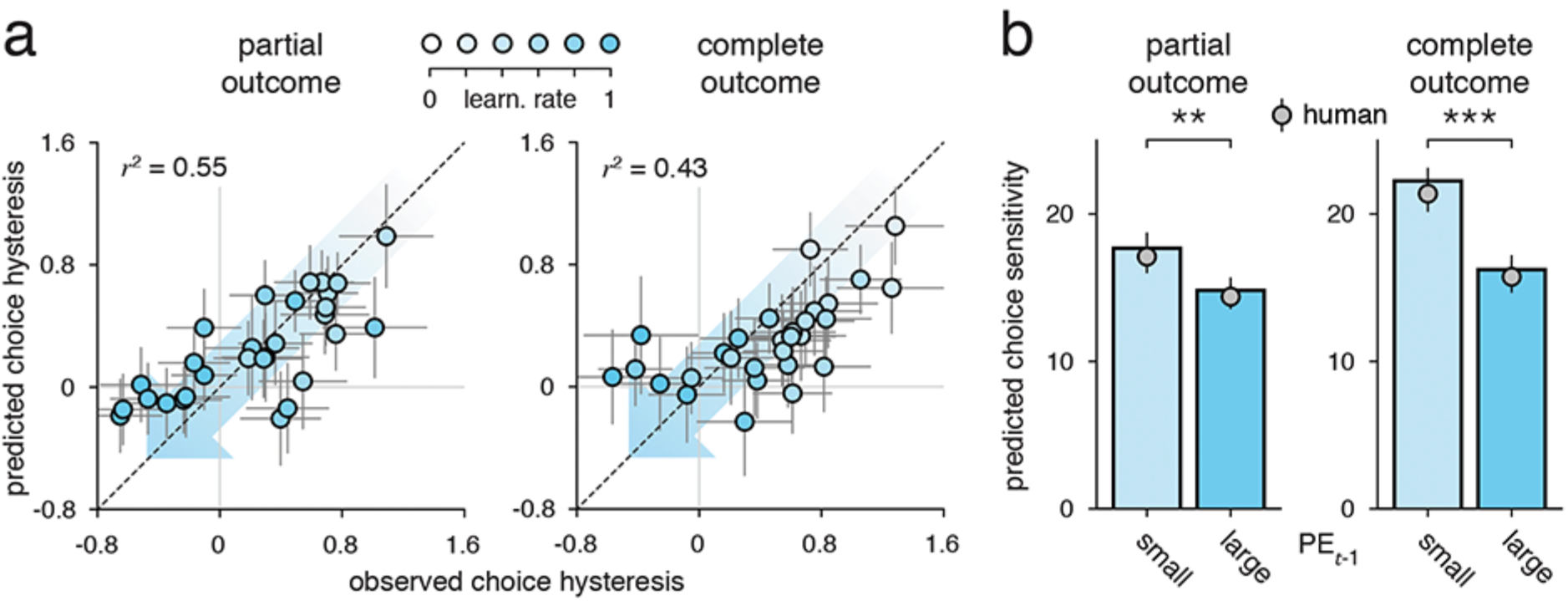
Characterization of decision effects predicted by learning noise. **a**) Relationship between the choice hysteresis measured in tested participants (x-axis) and the choice hysteresis predicted by simulations of the noisy RL model (y-axis) in the partial outcome condition (left panel) and the complete outcome condition (right panel). Dots ± error bars correspond to the mean ± s.d. of estimated posterior distributions for each tested participant. The blue shading of the dots indicates the learning rate associated with the chosen action (more saturated for faster learning rates). The choice hysteresis measured in tested participants correlates with the apparent choice hysteresis predicted by simulations of the noisy RL model in both conditions. Choice hysteresis also decreases with learning rate (shaded arrow). (**b**) Adaptation of choice stochasticity to surprise in tested participants (dots) and simulations of the noisy RL model (bars) in the partial outcome condition (left panel) and the complete outcome condition (right panel). The sensitivity of human choices fitted using an exact RL model (and measured through its best-fitting inverse temperature β) is significantly lower in trials following larger-than-average prediction errors, in both partial and complete outcome conditions. Simulations of the noisy RL model exhibit the same adaptation of choice sensitivity to surprise. Error bars correspond to s.e.m.

If choice hysteresis truly arises from the propagation of learning noise through action values, then it should naturally correlate with the degree to which learning noise propagates from one trial to the next, and thus decrease with learning rate. In other words, participants who learn more slowly should exhibit stronger choice hysteresis than participants who learn more rapidly. We tested this selective hypothesis in participants and obtained the predicted negative correlation between learning rate and choice hysteresis, in both conditions (Fig. 4a, linear correlation, partial: *r* squared = 0.691, d.f. = 27, *p* < 0.001; complete: *r* squared = 0.786, d.f. = 27, *p* < 0.001). Together, these findings indicate that the apparent choice hysteresis exhibited by participants can be parsimoniously explained as the consequence of noise in the underlying learning process.

A second effect documented in the literature consists in an adjustment of the softmax action selection policy to the surprise triggered by the preceding outcome – i.e., the magnitude of the prediction error in reinforcement learning^31–33^. Like choice hysteresis, this decision effect falls naturally out of learning noise, more specifically through its scaling *ζ* with the magnitude of the prediction error. Using an exact RL model to fit human decisions, we observed a decrease of the softmax inverse temperature *β* in trials following larger-than-average prediction errors, in both partial and complete outcome conditions (Fig. 4b, experiment 1, paired *t*-test, partial: *t*_28_ = −2.9, *p* = 0.007; complete: *t*_28_ = −3.7, *p* < 0.001). As for choice hysteresis, we then verified that simulations of the noisy RL model fitted using an exact RL model featured the same adaptation to surprise (paired *t*-test, partial: *t*_28_ = −5.5, *p* = 0.001; complete: *t*_28_ = −8.9, *p* < 0.001). Importantly, the adaptation predicted by simulations of the noisy RL model matched not only the direction, but also the size of the adjustment observed in participants (paired *t*-test, partial: *t*_28_ = 0.2, *p* = 0.847, BF_H0_ = 5.0; complete: *t*_28_ = 0.3, *p* = 0.778, BF_H0_ = 4.9). Furthermore, as predicted, participants with a steeper scaling *ζ* of learning noise with prediction errors showed a stronger adaptation to surprise (linear correlation, partial: *r* squared = 0.311, d.f. = 27, *p* = 0.002; complete: *r* squared = 0.313, d.f. = 27, *p* = 0.002). These results suggest that the apparent adaptation of choice stochasticity to surprise is caused by the multiplicative structure of noise in the underlying learning process, rather than by overt information seeking following surprising outcomes.

### Neural correlates of learning noise in the frontal cortex

Together, our quantitative dissection of non-greedy decisions points toward an important, yet previously unreported source of variability in reward-guided learning. To identify the neural mechanisms underlying this undocumented learning noise, we analyzed BOLD fMRI data (experiment 1, *N* = 29) recorded while participants performed the task (see Methods). We first used a standard model-free contrast (switch from *minus* repeat the previous action) to identify a classical network of brain regions implicated in cognitive control (Fig. 5a and Supplementary Table 1), including the dorsal anterior cingulate cortex (dACC), the dorsolateral prefrontal cortex (dlPFC), the frontopolar cortex (FPC) and the posterior parietal cortex (PPC). We then applied model-based analyses guided by the predictions of our noisy RL model to identify brain regions whose activations reflected the magnitude of learning noise corrupting each update of action values.

**Figure 5.**
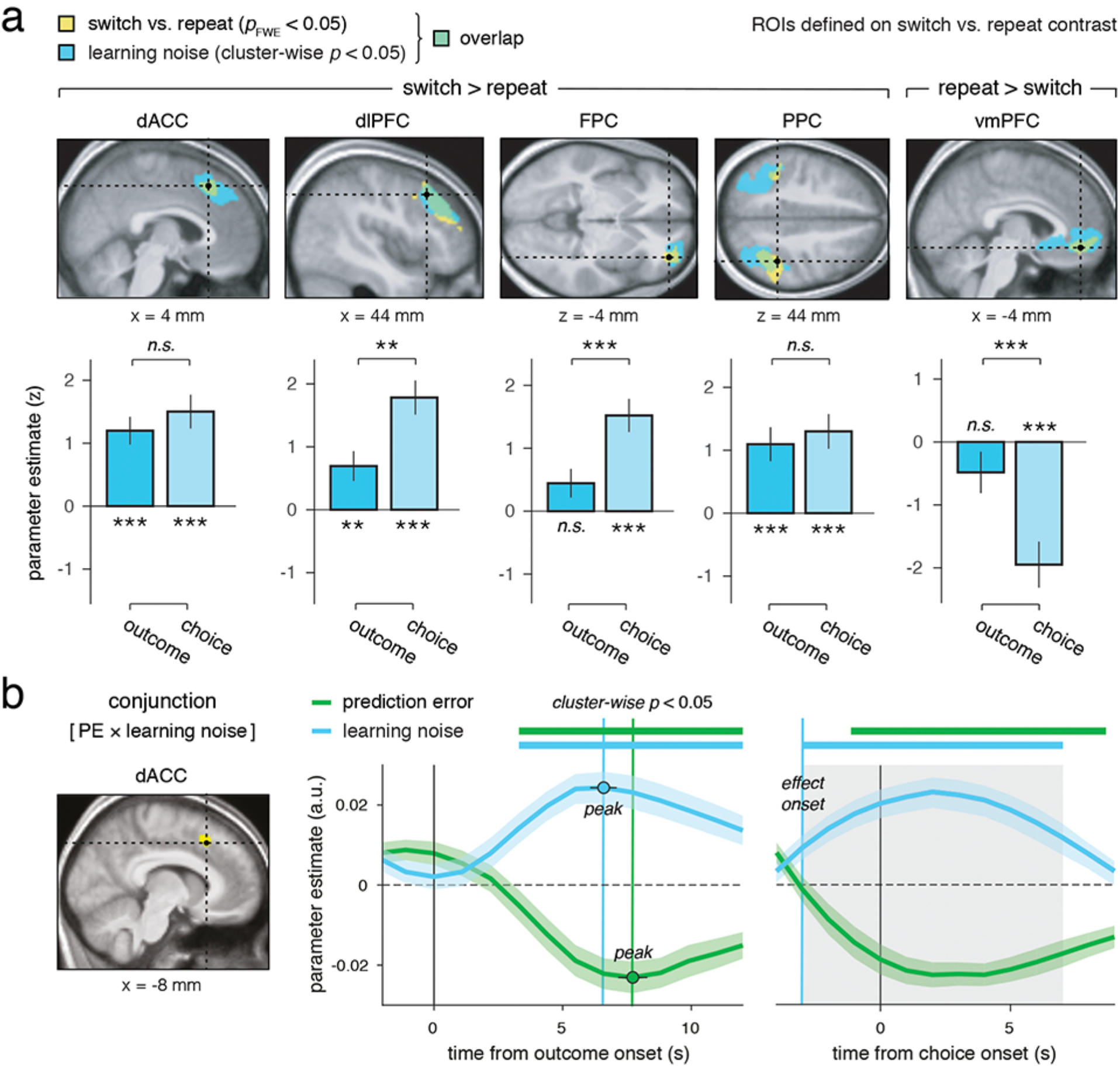
Neural correlates of learning noise in the human brain. (**a**) Upper panel: regions-of-interest (yellow) obtained for the switch minus repeat contrast corrected at a whole-brain family-wise error rate (FWE) of 0.05. Regions shaded in blue indicate the clusters that correlate significantly with learning noise at a cluster-wise corrected p-value of 0.05. Neural correlates of learning noise overlap broadly with neural correlates of the switch minus repeat contrast. Lower panel: group-level parameter estimates for the learning noise regressor in each of the ROIs defined by the switch minus repeat contrast locked to the onset of the outcome period of trial t−1 (left bar) and the choice period of trial t. Error bars correspond to s.e.m. Two stars correspond to a significant difference at p < 0.01, three stars at p < 0.001, n.s. to a non-significant difference. (**b**) Left panel: results of the whole-brain conjunction analysis of prediction error and learning noise at a cluster-wise corrected p-value of 0.05. Only the dACC reflects simultaneously the learning signal (i.e., the prediction error associated with the chosen action) and its trial-to-trial learning noise. Right panel: results of the finite impulse response (FIR) analysis showing the temporal dynamics of prediction error and learning noise in dACC activity, locked to outcome presentation (left) and to the onset of the following choice period (right). The correlations between dACC activity and prediction error (green) and learning noise (blue) peak at approximately the same time following outcome presentation (dots ± error bars correspond to jackknifed means and s.e.m. for correlation peaks estimated separately for the two regressors). Thick horizontal lines indicate time windows where parameter estimates diverge significantly from zero at a temporal cluster-wise corrected p-value of 0.01. The correlation between dACC and learning noise emerges significantly before the onset of the following choice period. Brain coordinates are expressed in the Montreal Neurological Institute (MNI) coordinate space.

For this purpose, we regressed canonical choice- and outcome-locked BOLD responses at each voxel against four trial-wise quantities derived from the noisy RL model fitted to each participant’s behavior: 1. the similarity between action values following each update step (choice ‘conflict’), 2. the difference between chosen and unchosen action values (choice ‘value’), 3. the prediction error associated with the obtained reward, and 4. the magnitude of learning noise corrupting the associated update of action values. This final quantity, specific to our noisy RL model, was computed as the predicted deviation |*ε*_*t*_| of noisy action values following each update step from the exact application of the Rescorla-Wagner rule to the same update step:

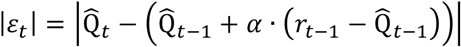

where 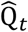 refers to noise-corrupted action values predicted by the noisy RL model conditioned on all observed rewards *r*_1:*n*_ and all actions *a*_1:*n*_ made by the participant. In practice, because summary statistics for 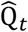 cannot be derived analytically, particle smoothing procedures were used to draw samples from the posterior distributions of 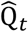, and |*ε*_*t*_| was averaged across drawn samples (see Methods). Importantly, the corresponding general linear model (GLM) was constructed using sequential orthogonalization to ensure that the noise regressor captured residual BOLD variance unaccounted for by the previous regressors also predicted by an exact RL model.

As documented in the literature, choice conflict was reflected positively in the dACC, whereas a broad network of brain regions correlated with choice value at a conservative statistical threshold (FWE-corrected *p* < 0.05) – with positive coefficients found in the ventromedial prefrontal cortex (vmPFC), and negative coefficients distributed across the dACC, the anterior insula, the dlPFC, the FPC and the PPC (Supplementary Table 2). The prediction error associated with the obtained reward (i.e., the learning signal used by the Rescorla-Wagner rule) was reflected positively in the ventral striatum and negatively in the dACC during the outcome period when action values are updated (FWE-corrected *p* < 0.05; Supplementary Table 2). Locked to the same event, an overlapping brain network reflected positively the magnitude of learning noise (i.e., trial-to-trial deviations from exact applications of the Rescorla-Wagner rule), including the dACC, the right dlPFC and the PPC (cluster-corrected *p* < 0.05; Fig. 5a). Importantly, a conjunction analysis revealed that among these three brain regions, only the dACC reflected simultaneously the prediction error and the magnitude of the associated learning noise during the outcome period when action values are updated (cluster-corrected *p* < 0.05; Fig. 5b).

To characterize more precisely the temporal dynamics of learning noise in dACC activity, we constructed a finite impulse response (FIR) model aligned either to the presentation of each outcome, or to the presentation of the following choice (see Methods). The positive correlation of dACC activity with the magnitude of learning noise peaked around the same time as the negative correlation with the prediction error following outcome presentation (Fig. 5c; learning noise: +6.1 s, prediction error: +7.7 s, jackknifed *t*_28_ = −1.7, *p* = 0.108). When aligned to the presentation of the following choice, the FIR model revealed that the correlation of dACC activity with learning noise emerged significantly before choice onset (Fig. 5b; jackknifed *t*_28_ = −2.7, *p* = 0.012). Together, these results indicate that dACC responses to obtained rewards reflect simultaneously the mean and variability of RL steps predicted by the Rescorla-Wagner rule.

During the following choice period when learning noise translates into behavioral variability, the magnitude of learning noise was reflected again positively in the dACC and the right dlPFC, but also in the FPC and negatively in the vmPFC (FWE-corrected *p* < 0.05; Fig. 5a). Importantly, the dACC reflected learning noise equally strongly in the outcome and following choice periods (dACC: *t*_28_ = 0.8, *p* = 0.390, BF_H0_ = 3.6). By contrast, the right dlPFC, the FPC and the vmPFC reflected learning noise significantly more strongly in the choice period (dlPFC *t*_28_ = 2.9, *p* = 0.008; FPC: *t*_28_ = 3.1, *p* = 0.005; vmPFC: *t*_28_ = −3.3, *p* = 0.003). Together, this pattern of findings assigns a unique position to the dACC among the neural correlates of learning noise: in addition to its correlation with the mean and variability of RL steps during outcome processing, dACC activity also reflects learning noise with the same intensity during the following choice period when learning noise triggers behavioral variability.

### Dissociating neural correlates of learning noise from surprise

Our noisy RL model hypothesizes that learning noise scales with the magnitude of the prediction error – i.e., the surprise associated with each outcome. And while dACC activity reflects the magnitude of learning noise in the outcome period, it has also been reported to monitor surprise in the same time period^34,35^. An important question is thus whether dACC activity reflects learning noise over and beyond its intrinsic correlation with surprise. Interestingly, the two quantities predicted by our noisy RL model shared only a third of variance with each other (linear correlation, *r* squared = 0.373 ± 0.086, mean ± s.e.m., ranging from 0.223 to 0.512 across participants).

To address this question, we first constructed two additional GLMs with sequential orthogonalization where we included surprise (i.e., the magnitude of the prediction error associated with each learning step) as an additional parametric regressor either before or after the learning noise regressor. Importantly, dACC activity was found to reflect learning noise over and beyond its intrinsic correlation with surprise (noise as last regressor, surprise: *t*_28_ = 3.6, *p* = 0.001; noise: *t*_28_ = 2.5, *p* = 0.019), whereas the converse was not true (surprise as last regressor, noise: *t*_28_ = 4.1, *p* < 0.001; surprise: *t*_28_ = −1.3, *p* = 0.195, BF_H0_ = 2.3). We then compared using neural BMS^22^ two additional GLMs where dACC activity was regressed against either the variability of learning steps (model 1) or the magnitude of prediction errors (model 2). Neural BMS revealed that model 1 provided a significantly better account of dACC activity than model 2 (exceedance *p* > 0.999). These findings indicate that, beyond their intrinsic correlation, dACC activity reflects learning noise rather than surprise: larger dACC responses to obtained rewards are associated with more variable, rather than larger, updates of action values. Note that a last GLM including both learning noise and surprise (model 3) provided a better fit than the two previous models (exceedance p > 0.999), suggesting that dACC activity reflects simultaneously learning noise and surprise.

### Dissociating neural correlates of learning noise from choice

We observed that learning noise is reflected in BOLD responses to obtained rewards when action values are updated, but also during the following choice period when learning noise triggers behavioral variability. This observation suggests that learning noise reflects not only neural variability in the update of action values, but also in the maintenance of chosen and unchosen action values until the next choice. However, this pattern may alternatively indicate that part of what is captured as learning noise by our noisy RL model truly arises from a non-modeled property of the choice process.

To address this important possible confound, experiment 1 included not only ‘choice’ trials where subjects could select the option they wanted to sample, but also ‘cued’ trials (Fig. 6a, 25% of all trials) where subjects were required to select one of the two shapes (pre-selected randomly by the computer, see Methods). In these cued trials, there is by definition no choice to be made, and indeed participants selected invariably the cued shape, in both partial and complete outcome conditions (Fig. 6b; partial: 98.3 ± 0.3%, complete: 96.6 ± 0.5%). We first verified that participants learnt equally from obtained rewards in choice and cued trials (learning rate *α*, choice: 0.603 ± 0.043, cued: 0.592 ± 0.039, *t*_28_ = 0.6, *p* = 0.581, BF_H0_ = 4.4). We then applied BMS to test whether learning in cued trials is corrupted by the same noise as in choice trials. As predicted by the noisy RL model, cued trials triggered significant learning noise (*ζ*_cued_ = *ζ*_choice_ vs. *ζ*_cued_ = 0, partial: BF ≈ 10^8.3^, exceedance *p* > 0.999; complete: BF ≈ 10^10.5^, exceedance *p* > 0.999).

**Figure 6.**
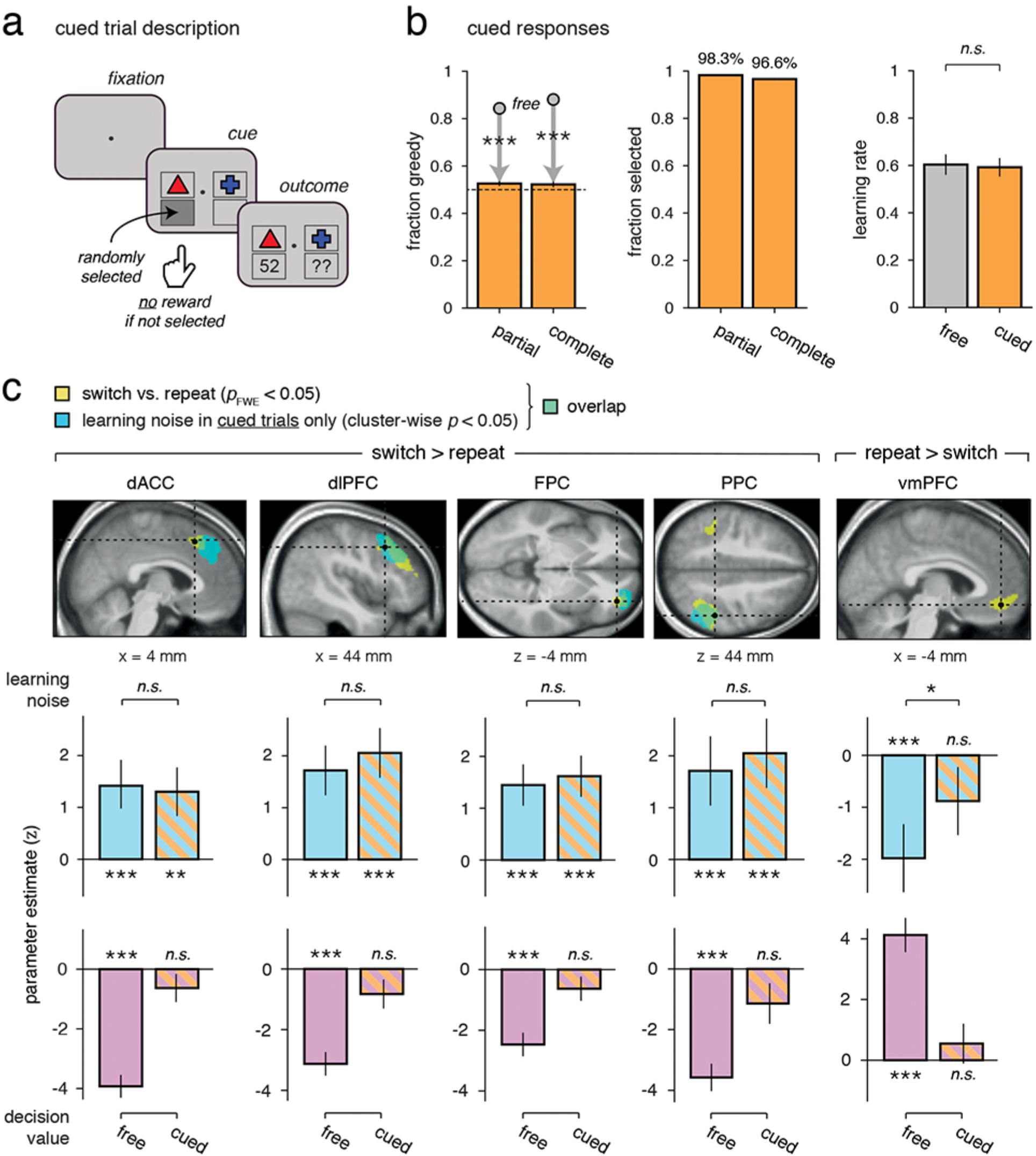
Neural correlates of learning noise in choice-free, cued trials. (**a**) Trial structure in ‘cued’ trials (1/4 of all trials). On cued trials, participants were required to select the highlighted action that was randomly pre-selected, and then observed its associated outcome (and the foregone outcome in the complete outcome condition) as in standard, ‘free’ trials. (**b**) Human behavior and learning in cued trials. Left panel: fraction of greedy (value-maximizing) actions in cued trials. As instructed, participants did not choose the highest-valued action in cued trials (dots indicate the fraction of greedy actions found in free trials as reference) both in the partial outcome condition (left bar) and the complete outcome condition (right bar). Middle panel: fraction of actions matching the pre-selected shape in cued trials. As instructed, participants almost invariably selected the highlighted action both in the partial outcome condition (left bar) and the complete outcome condition (right bar). Right panel: learning rates associated with the selected action estimated in free trials (left bar) and cued trials (right bar). Learning rates did not differ between free and cued trials, indicating that participants learnt equally from the two types of trials. (**c**) Upper panel: in contrast to Fig. 5, regions shaded in blue indicate the clusters that correlate significantly with learning noise in cued trials at a cluster-wise corrected p-value of 0.05. Neural correlates of learning noise remain significant in cued trials in all ROIs except the vmPFC. Lower panel: group-level parameter estimates for the learning noise regressor (above) and the decision value regressor (below) in each of the ROIs defined by the switch minus repeat in free trials (left bar) and cued trials (right bar). The decision value regressor is defined as the value difference between selected and unselected actions. The correlation of BOLD activity with decision value is significantly reduced in cued trials in all ROIs, whereas the correlation with learning noise is unchanged in all ROIs except the vmPFC. Error bars correspond to s.e.m. One star corresponds to a significant difference at p < 0.05, two stars at p < 0.01, three stars at p < 0.001, n.s. to a non-significant difference.

We could then make use of cued trials to test whether the neural correlates of learning noise found during the choice period were present even in cued trials. For this purpose, we constructed an additional GLM where choice and cued trials were modeled as separate events and modulated parametrically with: 1. the difference between selected and unselected action values, and 2. the magnitude of learning noise corrupting the preceding update of action values (see Supplementary Methods). The correlation of BOLD activity with the difference between selected and unselected action values in the dACC, the right dlPFC, the FPC and the vmPFC observed in choice trials was significantly reduced in cued trials where participants were required to select the cued shape (Fig. 6c; dACC: *t*_28_ = 5.2, *p* < 0.001; right dlPFC: *t*_28_ = 4.0, *p* < 0.001; FPC: *t*_28_ = 3.4, *p* = 0.002; vmPFC: *t*_28_ = 5.1, *p* < 0.001). No region remained significant at the whole-brain level in cued trials. These observations confirm that participants effectively did not choose the selected shape based on its action value in cued trials.

Nevertheless, and in agreement with our hypothesis, the positive correlation between BOLD activity and learning noise remained highly significant and unchanged in cued trials in the dACC (cued: *t*_28_ = 3.6, *p* = 0.001; cued vs. choice: *t*_28_ = −0.3, *p* = 0.760, BF_H0_ = 4.8), the right dlPFC (cued *t*_28_ = 6.0, *p* < 0.001; cued vs. choice *t*_28_ = 0.8, *p* = 0.404, BF_H0_ = 3.7) and the FPC (cued: *t*_28_ = 4.0, *p* < 0.001; cued vs. choice: *t*_28_ = −0.4, *p* = 0.677, BF_H0_ = 4.7). This finding further strengthens our hypothesis that the learning noise fitted by our noisy RL model reflects variability in the update of action values, rather than an unknown property of the choice process^3,7^. By contrast, the negative correlation between vmPFC activity and learning noise observed in choice trials was reduced in cued trials (cued: *t*_28_ = −1.6, *p* = 0.118, BF_H0_ = 1.6; cued vs. choice *t*_28_ = 2.8, *p* = 0.010), suggesting that its relationship with learning noise depends on the active selection of an action based on its expected value – the key task feature that is absent from cued trials.

### Relating neural correlates of learning noise to behavioral variability

Neuroimaging results so far indicate that BOLD activity in several brain regions (the dACC, the right dlPFC, the PPC, the FPC and the vmPFC) reflects learning noise corrupting each learning step. An important question arises here as to whether these different brain regions differ in their relationship with the resulting behavioral variability. To address this important question, we formulated a ‘brain-behavior’ analysis to predict behavioral variability based on trial-to-trial BOLD fluctuations in these five ROIs. In practice, we used a logistic regression model to predict each participant’s decisions to select either action as a function of the difference between action values, and single-trial deconvolved BOLD responses in the five ROIs (see Methods). The use of such ‘forward’ model of behavior, where BOLD responses in the different ROIs can be included simultaneously, affords to account for their shared variance. We reasoned that a neural signal reflecting learning noise should: 1. decrease participants’ sensitivity to the difference between action values (Fig. 7a), and 2. influence participants’ decisions also in the complete outcome condition where the choice process is best described by a purely value-maximizing action selection policy.

**Figure 7.**
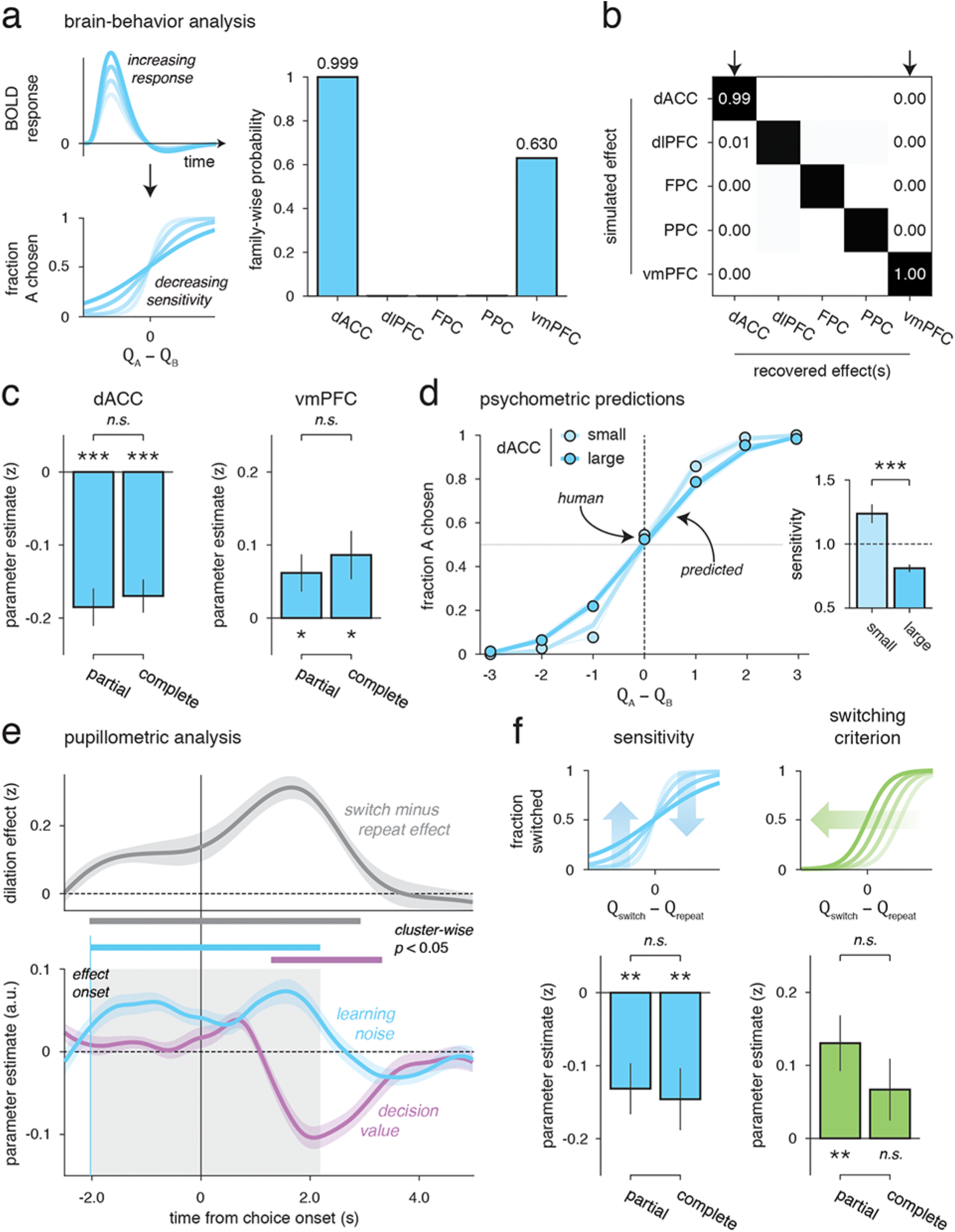
Brain-behavior and pupillometric analyses. (**a**) Left panel: schematic illustration of the brain-behavior relationship predicted by a neural correlate of learning noise. Trial-to-trial variability in the amplitude of BOLD responses (shaded in blue) should correlate negatively with trial-to-trial variability in the sensitivity to action values predicted by exact application of the Rescorla-Wagner rule on the previous trial. Right panel: results of the full factorial brain-behavior analysis including the five ROIs identified in Fig. 5. Family-wise probability is defined as the probability that each ROI modulates sensitivity independently of the involvement of other ROIs. Only the dACC and the vmPFC have family-wise probabilities exceeding 1%. (**b**) Model recovery results for the full factorial brain-behavior analysis. Confusion matrix displaying the estimated family-wise probabilities (columns) obtained for simulations of selective (single ROI) sensitivity modulations (rows). The full factorial brain-behavior analysis is able to recover accurately the source of simulated sensitivity modulations. (**c**) Participant-level parameter estimates for the winning model including the dACC (left panel) and the vmPFC (right panel) in the partial outcome condition (left bars) and the complete outcome condition (right bars). Sensitivity to action values correlates negatively with dACC activity and positively with vmPFC activity in both outcome conditions. (**d**) Psychometric brain-behavior predictions for the dACC ROI. Predicted (lines) and human (dots) psychometric curves for small (1^st^ tercile) and large (3^rd^ tercile) dACC responses. Both predicted and human curves show a decreased sensitivity to action values for larger dACC responses. Inset: sensitivity estimates for small (left bar) and large (right bar) dACC responses. Bars ± error bars correspond to jackknifed means and s.e.m. (**e**) Top panel: results of the switch minus repeat contrast for pupillary dilation. Pupillary dilation increases significantly before switches. Bottom panel: results of the finite impulse response (FIR) analysis showing the temporal dynamics of learning noise (blue) and decision value (purple) in pupillary dilation, locked to the onset of the choice period. The correlation between pupillary dilation and learning noise emerges significantly before the onset of the choice period. Thick horizontal lines indicate time windows where parameter estimates diverge significantly from zero at a temporal cluster-wise corrected p-value of 0.01. (**f**) Participant-level parameter estimates for pupil-linked modulations of sensitivity (left panel) and switching value/criterion (right panel) in the partial outcome condition (left bars) and the complete outcome condition (right bars). Pupillary dilation correlates negatively with sensitivity and positively with switching value. Error bars correspond to s.e.m. One star corresponds to a significant difference at p < 0.05, two stars at p < 0.01, three stars at p < 0.001, n.s. to a non-significant difference.

We entered single-trial BOLD responses *x*_*t*_ in the model as modulators of participants’ sensitivity to the difference between action values in the form of an interaction term 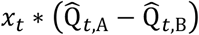. Importantly, 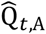 and 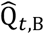 correspond to action values predicted by exact (noise-free) applications of the Rescorla-Wagner rule on the last update step, such that fluctuations in BOLD responses could be used to predict the behavioral effect of learning noise on the last update step. We followed a fully factorial scheme by constructing and estimating the posterior probabilities associated with all possible combinations of ROIs (32 = 2^5^, see Methods). We could then estimate the ‘familywise’ probability of each ROI to modulate sensitivity independently of the involvement of other ROIs. Note that, in contrast to posterior probabilities, these family-wise probabilities do not sum to one: they would be all equal to zero if no ROI modulates sensitivity, and all equal to one if all ROIs modulate sensitivity. This factorial analysis highlighted two ROIs with family-wise probabilities exceeding 1% (Fig. 7a,b): the dACC (family-wise *p* > 0.999) and the vmPFC (family-wise *p* = 0.630). Only two models (out of the 32 possible combinations) had posterior probabilities exceeding 1%: the winning model (posterior *p* = 0.629) including the dACC and the vmPFC, and the second-best model (posterior *p* = 0.370) including only the dACC.

Computing participant-level parameter estimates for the winning model (including the dACC and the vmPFC) in the partial outcome condition revealed a significantly negative effect of dACC responses on sensitivity (Fig. 7c; *β* = −0.185 ± 0.026, *t*_28_ = −7.2, *p* < 0.001), and a positive effect of vmPFC fluctuations on the same metric (Fig. 7c; *β* = 0.062 ± 0.026, *t*_28_ = 2.4, *p* = 0.023). The effect of dACC responses was substantially larger in absolute magnitude than that of vmPFC responses (+200.0%, *t*_28_ = 4.2, *p* < 0.001). In quantitative terms, these parameter values mean that sensitivity decreases by as much as 37% around its average value (standardized to one for each participant) when dACC responses increase by 2 standard deviations – hence conforming to the first signature of a neural signal reflecting learning noise (Fig. 7d). Furthermore, the estimated modulation of sensitivity by dACC responses was also significantly negative in the complete outcome condition (Fig. 7c; *β* = −0.170 ± 0.023, *t*_28_ = −7.4, *p* < 0.001). Importantly, the effects observed in the two conditions were similar (*t*_28_ = 0.5, *p* = 0.635, BF_H0_ = 4.6) – in line with the second signature of a neural signal reflecting learning noise. Together, these results support our hypothesis that dACC fluctuations reflect learning noise (which is present in both conditions) rather than choice stochasticity (which is absent in the complete outcome condition).

Finally, we tested whether the negative modulation of sensitivity by dACC responses could be caused not by *random* fluctuations of action values (as predicted by learning noise), but rather by *directed* fluctuations in the value of switching away from the previous action (as predicted by adjustments of the exploration-exploitation trade-off). To test this alternative brain-behavior relationship, we modified the logistic regression model to predict participants’ decisions to switch away from their previous action (i.e., *a*_*t*_ ≠ *a*_*t*-1_) as a function of the difference between the value of switching 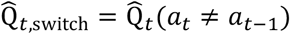 and the value of repeating the previous action 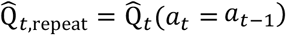. This time, we entered single-trial dACC responses *x*_*t*_ in the model not only as a modulator of participants’ sensitivity to the value difference between switching and repeating in the form of an interaction term 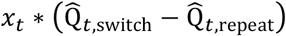, but also as a modulator of the value of switching in the form of an additive term *x*_*t*_. We followed a fully factorial scheme by constructing and estimating the model evidence associated with the 4 = 2^2^ combinations of the two possible modulations. This factorial analysis revealed a selective effect of dACC responses on sensitivity (family-wise *p* > 0.999), without any measurable effect on the value of switching (family-wise *p* < 0.001). In both partial and complete outcome conditions, the winning model included a selective effect of dACC responses on sensitivity (partial: posterior *p* = 0.963; complete: posterior *p* = 0.972). This selective brain-behavior relationship indicates that trial-to-trial fluctuations of dACC activity reflect learning noise rather than adjustments of the exploration-exploitation trade-off.

### Pupil-linked neuromodulatory correlates of learning noise

Beside frontal cortical contributions to non-greedy decisions, past research has identified the locus coeruleus-norepinephrine (LC-NE) system as a reliable neurophysiological correlate of behavioral variability^36–39^. Large phasic responses of LC neurons are associated with task disengagement and non-greedy decisions in particular^38,40^. While existing theories describe these effects as adjustments of the exploration-exploitation trade-off^33,37^, we hypothesized that trial-to-trial fluctuations of the computational precision of update steps, reflected in dACC activity, could be mediated by neuromodulatory fluctuations driven by the LC-NE system. Because LC activity is notoriously difficult to measure in fMRI, we took advantage of the strong, known correlation between LC activity and phasic pupil dilation^41–43^. We thus analyzed pupillary responses which were recorded in experiment 2 (*N* = 24 participants with clean data), by performing similar analyses to the ones conducted on BOLD signals in experiment 1 (see Methods).

As expected from the existing literature, we observed that a switch away from the previous action was associated with larger pupillary dilation in the preceding choice period and before (Fig. 7e; from −2.0 to 2.9 s following choice presentation, cluster-corrected *p* < 0.001). Pupillary dilation in the same time window correlated positively with the magnitude of learning noise |*ε*_*t*_| corrupting the preceding update step (Fig. 7e; from −2.0 to 2.2 s following choice presentation, cluster-corrected *p* < 0.001), and negatively with the value difference between chosen and unchosen actions (from 1.3 to 3.3 s following choice presentation, cluster-corrected *p* < 0.001). Interestingly, pupillary dilation started correlating with learning noise well before choice onset (*t*-test against zero, jackknifed *t*_23_ = −11.1, *p* < 0.001). This suggests that, like BOLD responses in the dACC, pupillary dilation reflects the variability of update steps during the learning of action values.

To confirm this hypothesis, we tested the relationship between trial-to-trial pupillary fluctuations and behavioral variability using the same brain-behavior analysis previously applied to BOLD responses in experiment 1. As for BOLD responses, we entered single-trial pupillary dilation in a logistic regression model of participants’ decisions as a modulator of their sensitivity to the difference between action values. The comparison between the ‘active’ model (where pupillary dilation predicts participants’ sensitivity) and the ‘null’ model (where pupillary dilation is not entered in the regression) indicated that pupillary dilation predicts participants’ sensitivity (posterior *p* > 0.999). Computing participant-level parameter estimates for this effect revealed a significantly negative effect of pupillary dilation on sensitivity (*β* = −0.198 ± 0.031, *t*_23_ = −6.3, *p* < 0.001). Like dACC responses, the estimated modulation of sensitivity by pupillary dilation was also significantly negative in the complete outcome condition (complete: *β* = −0.166 ± 0.036, *t*_28_ = −4.7, *p* < 0.001; complete vs. partial: *t*_28_ = 0.7, *p* = 0.520, BF_H0_ = 3.8).

As for BOLD responses, we then assessed how much this pupil-mediated reduction in sensitivity is caused by directed fluctuations in the value of switching rather than by random fluctuations in sensitivity (Fig. 7f). In the partial outcome condition, a factorial analysis indicated that pupillary dilation predicts both types of effects (random: family-wise *p* > 0.999; directed: family-wise *p* = 0.995). Computing participant-level parameter estimates for the two effects showed that increased pupillary dilation predicts simultaneously a lower sensitivity (Fig. 7f; *β* = −0.132 ± 0.035, *t*_23_ = −3.8, *p* = 0.001) and an increased value of switching (*β* = 0.130 ± 0.038, *t*_23_ = 3.4, *p* = 0.002). By contrast, in the complete outcome condition, increased pupillary dilation predicts random fluctuations in sensitivity (family-wise *p* > 0.999), but no directed fluctuations in the value of switching (familywise *p* = 0.095). Parameter estimation for each participant confirmed that increased pupillary dilation predicts a lower sensitivity (Fig. 7f; *β* = −0.146 ± 0.043, *t*_23_ = −3.4, *p* = 0.002), but no effect on the value of switching (*β* = 0.067 ± 0.042, *t*_23_ = 1.6, *p* = 0.129). Together, these results indicate that trial-to-trial fluctuations in pupillary dilation reflect learning noise over and above adjustments of the exploration-exploitation trade-off^44,45^.

## Discussion

Maximizing rewards in volatile environments requires an agent to trade the exploitation of currently best valued actions against the exploration of recently unchosen, and thus more uncertain ones. Dominant theories describe seemingly irrational, non-greedy decisions in terms of this exploration-exploitation trade-off, more specifically in terms of a drive to seek information about uncertain actions during choice^5,7,15,31,46,47^. Here we sought to contrast these information-seeking accounts with another possible source of behavioral variability: the limited computational precision of the learning process, which updates the expected values of possible actions following each reward. By decomposing behavioral variability into these two components using our noisy RL model, we show that more than half of non-greedy decisions are triggered by random noise in the learning process, rather than by an overt drive to seek information during choice.

This finding requires reconsidering the very nature of non-greedy decisions in volatile environments. Indeed, these decisions have classically been regarded as exploratory and information-seeking, and they should thus happen only (significantly more, at least) when there is uncertainty regarding the current value of recently unchosen actions. In accordance with this view, we found that the fraction of non-greedy decisions labeled as choice-driven by our noisy RL model depends critically on the absence of knowledge about the outcome of the foregone action on each trial. Our participants chose almost invariably the currently best-valued action when the outcome of the foregone action was observed or could be easily inferred from the outcome of the chosen action. By contrast, noise-driven variability in action values did not depend on knowledge about the foregone action, suggesting that its reflects a core characteristic of human learning rather than a feature that can be suppressed when the resulting behavioral variability is not useful^16^. In this sense, learning noise resembles the internal corruptive noise found in canonical decision-theoretic models, ranging from signal detection theory to sampling-based theories of inference^48–50^. Importantly, the predictions of our noisy model, cast in terms of reinforcement learning, are very similar to a noisy instantiation of ‘Kalman filtering’ (KF), which also provides an accurate description of human behavior in volatile environments^3,51^. Noise would be added in the KF model by assuming that the predictive means of action values are represented not by their full probability distributions, but a limited number of samples from these distributions. The moderate consistency of human decisions across identical blocks excluded the possibility that learning noise is due to a misspecification of our model – i.e., systematic deviations from the canonical Rescorla-Wagner rule used in our noisy RL model to update action values following each reward^17,52^.

The analysis of BOLD signals provided further information about the neural mechanisms underlying the observed learning noise. BOLD activity in a well-documented cognitive control network including the dACC and the FPC correlates positively with trial-to-trial deviations from the exact application of the canonical Rescorla-Wagner rule, even when participants were cued to select a randomly determined action – and thus did not have to make a choice. These neural correlates of learning noise are fundamentally different from neural correlates of computational quantities associated with reinforcement learning (e.g., prediction errors, expected values), in the sense that learning noise is neither computed explicitly by our noisy RL model nor hypothesized to be represented in any brain region. Our noisy RL model updates action values with a limited precision, and thus trial-to-trial deviations of each update step from the average reflect the effective variance (inverse precision) of the learning rule. This means notably that ‘functional’ accounts of surprise monitoring in the dACC might stem from the structure of learning noise^41,53–58^. In other words, dACC activity may correlate only indirectly with surprise through the multiplicative scaling of learning noise with the size of update steps – following the ubiquitous Weber’s law of intensity sensation found in numerous perceptual domains.

Several previous studies have tied both the dACC and the FPC to exploration and foraging across species^3,12,14,22,59–66^, but the specific contributions of these two regions have remained unclear. While both regions showed larger responses during switches, our brain-behavior analysis revealed that only dACC fluctuations exhibit the psychometric signatures of learning noise: a negative effect on participants’ sensitivity to action values, and a significant effect also in the complete outcome condition where there was no incentive to seek information about recently unchosen actions. By contrast, FPC fluctuations had no measurable effect on behavior in either condition. This relationship between dACC activity and learning noise has not been considered by previous studies and thus deserves further consideration. The dACC has been assigned a causal role in learning based notably on lesion studies in non-human primates disrupting reinforcement learning^67^, and on inactivation studies in rodents leading to purely random behavior divorced from learning^68^. The learning noise correlating positively with dACC activity is consistent with an active role of this frontal region in learning since the variability in question scales with the size of each learning step. Besides, several recent findings support the idea of learning-specific variability triggered by the dACC^69^. At the theoretical level, the ‘metaplastic’ synapses hypothesized in the dACC to account for adaptive learning in volatile environments go through stochastic transitions between states of faster and slower learning^70^. Neural circuits endowed with such synaptic properties would produce behavioral variability with the same statistical signatures as our learning noise. In the scanner, dACC activity has recently been shown to reflect prediction errors based on multiple, graded learning rates^69^. Neural variability in the pooling of such graded prediction errors (of which sampling would be an extreme case) would also produce behavioral variability of the same nature. Our findings reveal that the learning noise resulting from these accounts is responsible for a large fraction of non-greedy decisions.

An intriguing possibility is that learning noise may confer beneficial properties to the resulting behavior in volatile environments^16^. In particular, although the behavioral variability driven by learning noise does not have any active role in solving the exploration-exploitation trade-off, it provides a computationally inexpensive source of exploration by decreasing the precision of update steps^71^. Furthermore, as we have shown, the structure of learning noise has choice-stabilizing properties and produces an adjustment of exploration to surprise without requiring its explicit monitoring. Beyond these intrinsic benefits, computational noise in reinforcement learning may also optimize a second trade-off between the marginal payoff of a computation and the cost associated with performing the computation at a certain precision. The dACC has precisely been proposed to reflect a similar trade-off, by monitoring an ‘expected value of control’ (EVC) – defined as the difference between expected payoff and associated cost (cognitive conflict, in particular)^72^. Instead of assuming that the cost is represented explicitly in patterns of dACC activity, we propose that the cost associated with a computation may be reflected implicitly by its precision (Fig. 8a). This hypothesis provides a natural explanation as to why learning is subject to a limited computational precision, but also makes important testable predictions. In particular, increasing the level of volatility (i.e., the rate of change in the average values of actions) reduces the marginal payoff of learning (which in the limit case tends toward zero), and thus decreases the precision which optimizes the underlying payoff-cost trade-off (Fig. 8b). We thus predict that participants should feature not only larger learning rates^9,73^, but also more learning noise, in more volatile environments. Under our hypothesis, the larger dACC activity observed at higher levels of volatility^9^ would be a signature of the increased learning noise predicted in such conditions.

**Figure 8.**
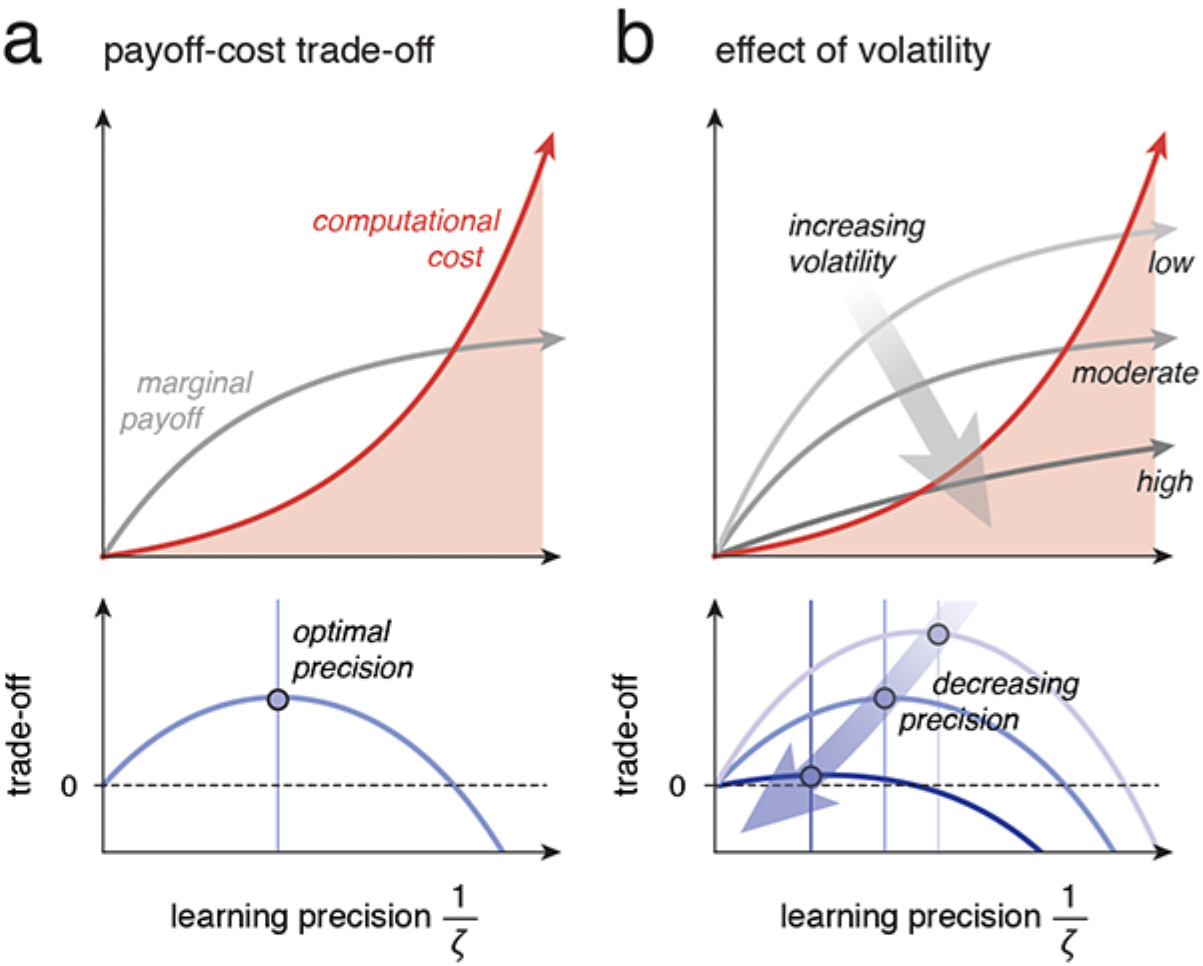
Hypothesized payoff-cost trade-off on learning precision. (**a**) Illustration of the hypothesized payoff-cost trade-off. Top panel: schematic illustration of the dependencies between learning precision 1/ζ (x-axis) and the marginal payoff of learning (gray curve) and the computational cost of learning (red curve). The marginal payoff of learning saturates at high precisions, whereas its computational cost grows exponentially. Bottom panel: payoff minus cost trade-off showing an optimal learning precision which maximizes the difference between payoff and cost. (**b**) Illustration of the effect of volatility on the hypothesized payoff-cost trade-off. Top panel: increasing volatility decreases the marginal payoff of learning (from lighter to darker gray curves). Bottom panel: increasing volatility decreases the optimal learning precision which maximizes the difference between payoff and cost (from lighter to darker blue curves).

Based on previous findings, we reasoned that the observed learning noise, reflected in dACC responses, may be linked to the ongoing state of the locus coeruleus-norepinephrine (LC-NE) system – which has been previously involved in both the regulation of the neural gain of cognitive operations and the adjustment of the exploration-exploitation trade-off^32,33,36–38,40^. Indeed, LC neurons receive strong projections from the dACC, which in turn produce gain control in several frontal regions implicated in reward-guided learning^74–76^. Pupil-linked fluctuations of the LC-NE state are associated with both task disengagement and non-greedy decisions^39,77,78^. Existing theories have interpreted these findings as evidence in favor of the implication of the LC-NE system in controlling the exploration-exploitation trade-off, something for which there is only partially conclusive evidence to date^37,39,77^. We hypothesized that the LC-NE system may instead mediate the relationship between dACC activity and learning noise. In line with this hypothesis, we observed that pupillary dilation predicts not only directed fluctuations in the value of switching away from the previous action, but also random fluctuations in participants’ sensitivity to action values. This was the case even when participants observed the outcome of the foregone action on every trial, just like dACC fluctuations. This relationship between pupillary dilation and sensitivity supports the idea that the LC-NE system controls the proposed payoff-cost trade-off by adjusting the computational precision of learning. The recent observation that pupillary dilation increases at high levels of volatility^73^ (i.e., when the optimal learning precision decreases) provides preliminary evidence for this hypothesis, which should be tested formally in future work.

Together, our findings emphasize a large, yet previously neglected source of random variability in reward-guided decision-making, driven by computational noise in the underlying learning process. This noise-driven source of non-greedy decisions, likely occurring unbeknownst to the decision-maker, is independent of the overt arbitration between the exploitation of currently best-valued actions against the exploration of more uncertain ones – a trade-off previously considered as the sole source of behavioral variability. As we have shown, the decomposition of non-greedy decisions into noise- and choice-driven components bears important consequences for understanding both the mechanisms underlying reward-guided behavior and its neurophysiological substrates. Exact models of learning should be revised to allow for noise in their core computations, and include an explicit cost-benefit trade-off regulating their precision.

## Methods

### Participants

fMRI experiment 1: 30 subjects (16F, mean age 26.0 ± 5.5, all right handed). One subject was excluded from the analysis because he failed to understand the task instructions and performed at a chance level. Subjects were screened for the absence of any history of neurological and psychiatric disease or any current psychiatric medication and had normal or corrected to normal vision.

Behavioral experiment 2: 30 subjects (17F, 23.6 ± 4.6) took part in the experiment and all the data were included in the analysis. Additionally to the behavioral data, we also recorded the pupillary responses in this study. However, N = 6 subjects were excluded from pupillatory analyses because of bad quality pupil data.

Behavioral experiment 3 (with the repeated blocks): 30 subjects (19F, mean age 24.2 ± 3.7) took part in the experiment and all the data were included in the analysis.

All participants in all three experiments gave a written informed consent and the local Ethical Committee approved the study. All subjects received a fixed payment and were additionally remunerated based on their performance in the task between 5 and 10 euros.

### Experimental task

In the three experiments, we asked subjects to play a two-armed bandit task where the reward magnitudes of the two bandits followed a random walk process (Fig. 1a). The rewards observed by the subjects (between 1 and 99 points) were sampled from Gaussian distributions with mean predicted by the random walks (Fig. 1b). In experiment 1 (fMRI study) and 2, in half of the experimental blocks (4 out of 8), the reward obtained on each trial from the chosen lever was presented simultaneously with the ‘counterfactual’ feedback that could have been obtained from the unchosen lever - referred to as the complete setting [10], while, in the other half, only the chosen outcome was shown - referred to as the partial setting. In experiment 3, subjects were always presented with the full information about both obtained and forgone outcomes but in half of the blocks unbeknownst to the participants the reward sequences were repeated (Fig. 3a).

### Analytical Model

The full analytical model of behavior is:

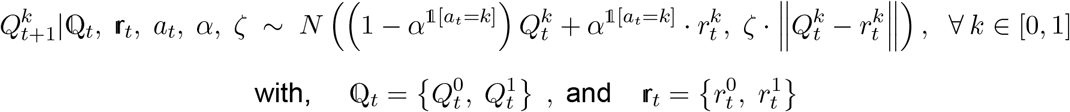

With 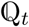 the Q-values at time t, *α* the learning rate, *ζ* the learning noise scaling and 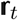 the observed or fictive rewards at time t. For the complete setting, 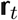 are the two observed rewards. For the partial setting, the regression to the mean models set one of these rewards to their empirical mean - 50. The dynamics of the actions are :

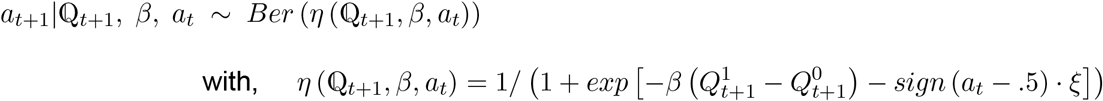

With *β* the softmax coefficient and *ξ* the repetition bias. Let us add a parameter *c* indicating whether the two options have a shared (c=1) or different (c=0) learning rate. Working in a Bayesian framework, we ascribe a prior distribution to each parameter:

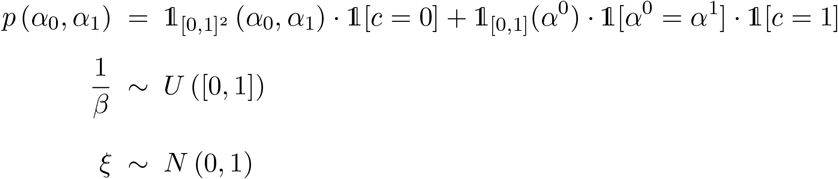

All the models can be derived from these equations by setting parameters to 0. For instance *ζ* and *ξ* to 0 leads to the standard reinforcement learning model.

The standard deviation of the learning noise is proportional to the prediction error 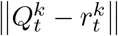 which implies it scales positively with the quantity of update 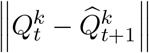, with 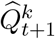 obtained by applying the exact reinforcement learning update to 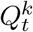.

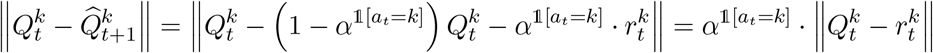

### Fitting procedure

Fits in all models are based on Monte Carlo methods [11]. More precisely, for the standard reinforcement learning models without learning noise, we used an iterated batch importance sampler (IBIS) [1]. IBIS is a sequential Monte Carlo (SMC) algorithm for exploring a sequence of parameter posterior distributions when the likelihoods *p*(*a*_*t*_|*a*_1:(*t*−1)_*, r*_1:(*t*−1)_*, β, α*^0^*, α*^1^*, ξ*) are tractable. This algorithm could not be used for the models with learning noise as these likelihoods are, in this latter case, intractable. Thus, we used the SMC^2^ algorithm [2] to perform inference in the models with learning noise. These fitting procedures can be found in more extensive details in the Supplementary Information.

### Obtaining the smoothing distributions

Two studies involving the learning noise model required the smoothing distributions 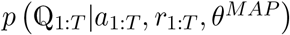, with *θ*^*MAP*^ the maximum a posteriori. The first was when we investigated whether a labeled ‘non-greedy’ trial effectively stemmed from the softmax contribution or whether it was a greedy choice that originated from learning noise. The second study involving the smoothing distributions is the model-based fMRI study : we correlated the BOLD signal with the most likely deviations predicted by the learning noise model. To obtain samples approximately distributed under the smoothing distributions, we used the Forward Filter/Backward Simulator (FFBSi) [5] [7] to obtain N samples 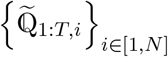 from 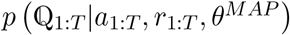 with T the total number of trials.

### Distinguishing softmax stochasticity from learning noise

In the main experiment and its purely behavioral extension, we reported the proportion of labeled ‘non-greedy’ trials which originated from learning noise. To label a trial as ‘non-greedy’, we applied the standard exact reinforcement learning algorithm and labeled as ‘non-greedy’ every trial which was not predicted by this exact model [3]. Assuming there are a total of T trials and among these T trials, n were labeled as ‘non-greedy’.

Let us write {*t*_1_, …, *t*_*n*_} the indexes of these n trials. For each *t*_*i*_, given the smoothing average 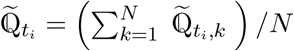, the ‘non-greedy’ trial *t*_*i*_ can actually be greedy for the learning noise model if it is well predicted by the sign of 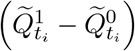. We conclude the proportion of ‘non-greedy’ trials to be greedy for the learning noise model to be:

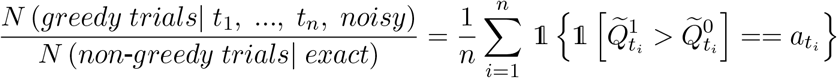

### Obtaining the learning noise and softmax variabilities

For the computation of the learning noise variability, please refer to the paragraph ‘Obtaining the noise regressors for model-based fMRI’ : for each subject, we obtained the learning noise variability by calculating the variance over the deviations *δ*_1:*T*_. As for the computation of the softmax variability, we used the approximation of the logistic distribution as the cumulative distribution function of a Gaussian distribution (see Supplementary Information). From this approximation, we obtained the stochasticity added by the logistic distribution with softmax parameter *β* was well-approximated by a Gaussian noise with standard deviation 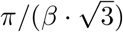

### Obtaining the bias/variance tradeoff

The last experiment (exp.3) composed of repeated blocks was used to obtain a bias/variance tradeoff describing to which extent the learning noise can be explained by deterministic deviations from the learning rule. This experiment led to a consistency ratio of 82% indicating that, on 82% of the trials, the subjects were consistent in their choices within repeated blocks. For each set of fitted parameters (one per subject with N=30 subjects), we simulated the model with learning noise and no softmax stochasticity a 100 times. This led to 30 × 100 agents playing the task. Furthermore, in these simulations, we assumed a shared contribution throughout the repeated blocks and a unique contribution to each of them. The shared contribution represented the deterministic deviation to the learning rule, the “bias”, whereas the unique contributions represented unpredictable deviations, the “variance”. We modulated the bias/variance contributions so as to match the consistency ratio of the subjects and obtained contributions of 68% of variance and 32% of bias. This means the learning noise obtained is only explained up to one third by deterministic biases (fig 3d).

### Obtaining the noise regressors for model-based fMRI

For the model-based fMRI, we investigated the fMRI BOLD activity correlates with regressors issued from the model with learning noise. With *T* the total number of samples and *N* the number of particle, let 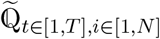 be the smoothing samples of the learning noise model and 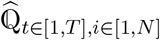 the Q-values obtained when applying the exact update rule to 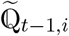 (see paragraph ‘Obtaining the smoothing distributions’). The fMRI regressors include the prediction error of the chosen option 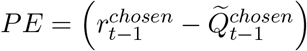, with 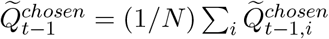, the difficulty 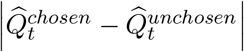, with 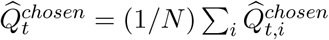 and 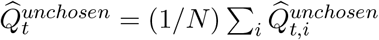, and the deviations from the exact Rescorla-Wagner learning rule at every time step. This last ‘learning noise’ regressor is obtained by computing the deviation for each sampled trajectory, 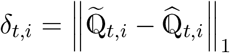 and averaging over these trajectories *δ*_*t*_ = (1/*N*)∑_*i*_ *δ*_*t,i*_ (main text equation 3).

### Computing the Bayes factor of the null hypothesis

When standard t-tests were considered non-significant, we performed an additional Bayesian test to obtain the Bayes factor quantifying the evidence in favor of the null hypothesis. This additional test was computed using the ‘BayesFactor’ library under R implementing the test described in Rouder JN. and colleagues, 2009 [12].

### Imaging data acquisitions and preprocessing

A Siemens Prisma fit 3T scanner (CENIR, ICM, Paris, France) and an 64-channel head coil were used to acquire both high resolution T1-weighted anatomical MRI using a 3D MPRAGE with a resolution of 1*mm*^3^ voxel and *T* 2*-weighted multibandecho planar imaging (mb-EPI) with multiband factor of 3 and acceleration factor of 2 (GRAPPA). The parameters of the fMRI time series acquisition were the following: 54 slices acquired in ascending order, the voxel size was 2.5mm isometric (in each direction), the repetition time of 1.1s, and the echo time of 25 ms. A tilted plane acquisition sequence was used to optimize sensitivity to BOLD signal in the orbitofrontal cortex [4], [13]. Preprocessing included co-registration of the anatomical T1 images with the mean EPI, segmentation and normalization to a standard T1 template, and average across all subjects to allow group-level anatomical localisation.

Preprocessing of the mb-EPI consisted of spatial realignment, movement correction, reconstruction and distortion correction, and normalization using the same transformation as applied for the structural images. Normalized images were spatially smoothed using a Gaussian kernel with a full width at a half-maximum of 8mm. All the preprocessing except for the distortion correction was done using the SPM12 (Wellcome Trust Center for NeuroImaging, London, UK; www.fil.ion.ucl.ac.uk). Distortion correction consisted of image unwarping and reconstruction done using FSL software [6].

### FIR time courses analysis

To reconstruct the time courses of BOLD signal showed in Fig. 5b, we created two GLMs. For the first GLM, the time epoch started at 2 seconds before up to 12 seconds after the onset of outcome and then applied a GLM to each timepoint separately. The GLM1 included the following parametric modulators: reward prediction error for the chosen option and the learning noise regressor, sequentially orthogonalized as in the other GLMs. For the GLM 2 we chose a time epoch starting at 3 seconds before up to 10 seconds after the onset of choice. The GLM 2 included the following parametric modulators: the reward prediction error for the chosen option from the previous trial and the learning noise regressor for the current trial. Obtained time courses for parametric modulators in both GLMs were first smoothed using Savitzky-Golay filtering (order 2, length 7 timepoints) and then averaged across subjects. For GLM 1, we next computed the time to peak for each parametric modulator using leave-one-out procedure and then averaged the estimates across subjects. Individual time-to-peak estimates were brought to a paired two-tailed t-test with the p-value adjusted for the resampling (Jacknife resampling method, [9]). For both GLMs, we determined the time window where parametric modulators for learning noise, prediction error and relative choice value were significantly different from zero. We used non-parametric cluster correction (p < 0.01) using permutation method (N = 10.000) on the obtained regression coefficients [8].

### Predicting the behavioral variability using the fMRI time-series data

We ran logistic regression analysis to determine the role of each of the five ROIs (dACC, dlPFC, PPC, FPC and vmPFC) which were shown to reflect the learning noise in the modulation of decision sensitivity (Fig. 7a). First, we estimated the individual sensitivity to the relative value difference for each subject. To do this, we ran a simple logistic regression model: *P* (*choose A*) = *probit*(*β*_0_ + *β*_1_ · (*Q*^*A*^ − *Q*^*B*^)) to compute the adjusted value difference, ∆*Q*_*adj*_ = *β*_0_ + *β*_1_ · (*Q*^*A*^ − *Q*^*B*^), and then used this adjusted value difference as a predictor in the main logistic regression analyses. This adjustment allowed us to account for individual differences in decision sensitivity and to quantify the contribution of ROIs in terms of effect size to the behavioral variability. We next predicted the trial-by-trial decision to choose option A as a function of the adjusted difference between two options values ∆*Q*_*adj*_ and z-scored trial-by-trial deconvolved BOLD signal as a modulator of participant’s sensitivity to the relative option value (e.g, for the dACC, ∆*Q*_*adj*_ · *dACC*). For 5 ROIs we constructed 2^5^ regression models to estimate the model evidence associated with all possible combinations of ROIs. Each model was fitted across all subjects simultaneously (fixed-effect analysis). For each region independently we summed up model evidence for all models that included this region to compute the ‘family-wise’ probability (Fig. 7a). The winning model based on the family-wise analysis included the following terms: *P* (*choose A*) = *probit*(*β*_1_ · ∆*Q*_*adj*_ + *β*_2_ · *dACC* · ∆*Q*_*adj*_ + *β*_3_ · *vmP FC* · ∆*Q*_*adj*_). We next fitted this model separately for each subject in the partial and complete feedback conditions and next extracted the regression estimates for the two ROI modulatory effects to perform two-tailed paired t-tests (see Results).

### Predicting decisions using the trial-to-trial pupil dilation

Pupil data were acquired in a separate dataset (experiment 2, N = 30 with pupil data successfully recorded for 24 subjects) who performed the same task outside the scanner. The diameter of the dominant eye was recorded at 500 Hz using an EyeLink-1000 System (EYELINK II CL v4.594). Subjects performed the experiment in a dark sound-proof room with their head positioned on a chin rest positioned at approximately 50 cm from the computer screen.

Data were first corrected for the eye-blinked artifacts using manufacturer default algorithm. The preprocessing was then performed using custom-based MatLab functions: data were smoothed using the moving average of of 100 ms; blink periods were linearly interpolated (from −100 ms before until 500 ms after the blink). Trials where blinking or fixation errors occurred within the analysis window (from −1000 ms till 5000 ms after the stimulus onset) were removed. Data were next subsampled at 50 Hz and low-pass filtered at 4Hz (third order Butterworth filter) and then z-scored per session.

We next analyzed these time series using FIR approach. First, we check that pupillary response increases after the switch vs. repeat decisions: the FIR model across both partial and complete feedback conditions included stick function put at every time point within interval 1000 ms before and 5000 ms after the stimuli onset and then parametrically modulated with the switch (1) vs. repeat (−1) regressor. We next constructed another FIR model within the same time window where each stick function was parametrically modulated with the two computational variables: the relative value (Qchosen - Qunchosen) at the time of choice t obtained by applying the exact update to the noisy Q-value and the learning noise. Recalling the notations of the ‘obtaining the noise regressors for model-based fMRI’ paragraph, these two regressors correspond to 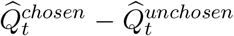 and *δ*_*t*_. All behavioral variables were sequentially orthogonalized as for the fMRI analysis, z-scored per session; additional session regressors were added to the model. The model was next inverted using the Ridge regression. For both FIR models, we performed nonparametric cluster correction on the obtained regression coefficients as described in Maris and Oostenveld [8]. Clusters were first defined for 8 minimum contiguous time points where the beta estimates were different from 0 at p < 0.01 for two-tailed paired t-test and the summed t-statistic was computed for each cluster. Next permutation analysis was run (N = 10.000) where each time half of the subjects betas were flipped in sign and the sum of the t-statistic values was recomputed. Significant cluster was defined at p < 0.01 (1% chance that the shuffled t-value exceeded the true summed t-value for a given cluster).

We defined a significant time interval where the pupil dilation negatively correlated with the relative value in the partial and complete feedback conditions (Fig. 7e). We next averaged trial-by-trial pupil response within this time interval and used these averaged response in the logistic regression model to predict subjects’ with decision to choose option A on each trial. The logistic regression was similar to the one used to investigate the effect of BOLD fluctuations with the frontal regions on subjects’ behavior and consisted of the following regressors: subject-by-subject adjusted relative option value ∆*Q*_*adj*_ (see previous paragraph) at the time of choice t without the added noise on this trial from the noisy computational model and the trial-by-trial interaction between the z-scored pupil fluctuations and the adjusted relative option value, *P* (*choose A*) = *probit*(*β*_1_ · ∆*Q*_*adj*_ + *β*_2_ · *pupil* · ∆*Q*_*adj*_). Individual regressor estimates were next brought to a second-level between-subject analysis using paired t-tests.

### Code availability

Python and C++ code for the fitted exact and noisy RL models are available at https://github.com/csmfindling/learning_variability and the principal algorithms can be found in the Supplementary Information.

### Data availability

The data that support these findings are available from authors upon request.

## Acknowledgments

We thank C. Summerfield for comments on an earlier version of the manuscript. This work was supported by a starting grant from the European Research Council awarded to V.W. (ERC-StG-759341), a junior researcher grant from the Agence Nationale de la Recherche awarded to V.W. (ANR-14-CE13-0028), and two department-wide grants from the Agence Nationale de la Recherche (ANR-10-LABX-0087 and ANR-10-IDEX-0001-02 PSL). C.F. was supported by a graduate research fellowship from the Direction Générale de l’Armement (2015-60-0041). S.P. was supported by a CNRS-Inserm ATIP-Avenir grant (R16069JS) and a research grant from the Programme Emergence(s) of the City of Paris.

Author contributions
Conceptualization: S.P. and V.W.; Methodology: C.F., V.W., S.P., and V.W.; Formal Analysis: C.F., V.S., and V.W.; Investigation: V.S. and R.D.; Writing – Original Draft: C.F., V.S., and V.W.; Writing – Review & Editing: C.F., V.S., S.P., and V.W.; Supervision: V.W.; Funding Acquisition: V.W.

## References

1. Sutton, R. S. & Barto, A. G. Reinforcement learning: an introduction. MIT Press, Cambridge, MA (1998).

2. Rescorla, R. A. & Wagner, A. R. A theory of Pavlovian conditioning: variations in the effectiveness of reinforcement and nonreinforcement. Classical Conditioning II, eds. Black, A. H. & Prokasy, W. F., 64–99. Appleton-Century-Crofts, New York, NY (1972).

3. Daw, N. D., O’Doherty, J. P., Dayan, P., Seymour, B. & Dolan, R. J. Cortical substrates for exploratory decisions in humans. Nature 441, 876–9 (2006).

4. Lee, M. D., Zhang, S., Munro, M. & Steyvers, M. Psychological models of human and optimal performance in bandit problems. Cogn. Syst. Res. 12, 164–174 (2011).

5. Frank, M. J., Doll, B. B., Oas-Terpstra, J. & Moreno, F. Prefrontal and striatal dopaminergic genes predict individual differences in exploration and exploitation. Nat. Neurosci. 12, 1062–1068 (2009).

6. Zhang, S., Huang, C. H. & Yu, A. J. Sequential effects : a Bayesian analysis of prior bias on reaction time and behavioral choice. Proc. Annu. Meet. Cogn. Sci. Soc. 1844–1849 (2014).

7. Wilson, R. C., Geana, A., White, J. M., Ludvig, E. A. & Cohen, J. D. Humans use directed and random exploration to solve the explore–exploit dilemma. J. Exp. Psychol. Gen. 143, 2074–2081 (2014).

8. Kolling, N., Behrens, T. E. J., Mars, R. B. & Rushworth, M. F. S. Neural mechanisms of foraging. Science 336, 95–8 (2012).

9. Behrens, T. E. J., Woolrich, M. W., Walton, M. E. & Rushworth, M. F. S. Learning the value of information in an uncertain world. Nat. Neurosci. 10, 1214–21 (2007).

10. Rushworth, M. F. S. & Behrens, T. E. J. Choice, uncertainty and value in prefrontal and cingulate cortex. Nat. Neurosci. 11, 389–97 (2008).

11. Shenhav, A., Straccia, M. A., Cohen, J. D. & Botvinick, M. M. Anterior cingulate engagement in a foraging context reflects choice difficulty, not foraging value. Nat. Neurosci. 17, 1249–54 (2014).

12. Boorman, E. D., Behrens, T. E. & Rushworth, M. F. Counterfactual choice and learning in a neural network centered on human lateral frontopolar cortex. PLOS Biol. 9, e1001093 (2011).

13. Koechlin, E. & Hyafil, A. Anterior prefrontal function and the limits of human decision-making. Science 318, 594–598 (2007).

14. Donoso, M., Collins, A. G. E. & Koechlin, E. Foundations of human reasoning in the prefrontal cortex. Science 344, 1481–6 (2014).

15. Gershman, S. J. Deconstructing the human algorithms for exploration. Cognition 173, 34–42 (2018).

16. Wyart, V. & Koechlin, E. Choice variability and suboptimality in uncertain environments. Curr. Opin. Behav. Sci. 11, 109–115 (2016).

17. Drugowitsch, J., Wyart, V., Devauchelle, A.-D. & Koechlin, E. Computational precision of mental inference as critical source of human choice suboptimality. Neuron 92, 1398–411 (2016).

18. van der Helm, P. A. Weber-Fechner behavior in symmetry perception? Attention, Perception, Psychophys. 72, 1854–1864 (2010).

19. Fechner, G. T. Elements of psychophysics, trans. Adler, H. Holt, Reinehart & Winston, New York, NY (1966).

20. Krueger, L. E. Perceived numerosity: a comparison of magnitude production, magnitude estimation, and discrimination judgments. Percept. Psychophys. 35, 536–542 (1984).

21. Palminteri, S., Wyart, V. & Koechlin, E. The importance of falsification in computational cognitive modeling. Trends Cogn. Sci. 21, 425–433 (2017).

22. Palminteri, S., Khamassi, M., Joffily, M. & Coricelli, G. Contextual modulation of value signals in reward and punishment learning. Nat. Commun. 6, 8096 (2015).

23. Klein, T. A., Ullsperger, M. & Jocham, G. Learning relative values in the striatum induces violations of normative decision making. Nat. Commun. 8, 1–12 (2017).

24. Li, J. & Daw, L. Signals in human striatum are more appropriate for policy update rather than value prediction. J. Neurosci. 31, 5504–5511 (2011).

25. Lau, B. & Glimcher, P. W. Dynamic response-by-response models of matching behavior in rhesus monkeys. J. Exp. Anal. Behav. 84, 555–579 (2005).

26. Schönberg, T., Daw, N. D., Joel, D. & O’Doherty, J. P. Reinforcement learning signals in the human striatum distinguish learners from nonlearners during reward-based decision making. J. Neurosci. 27, 12860–7 (2007).

27. Gershman, S. J., Pesaran, B. & Daw, N. D. Human reinforcement learning subdivides structured action spaces by learning effector-specific values. J. Neurosci. 29, 13524–31 (2009).

28. Padoa-Schioppa, C. Neuronal origins of choice variability in economic decisions. Neuron 80, 1322–1336 (2013).

29. Kiyonaga, A., Scimeca, J. M., Bliss, D. P. & Whitney, D. Serial dependence across perception, attention, and memory. Trends Cogn. Sci. 21, 493–497 (2017).

30. Yu, A. J. & Cohen, J. D. Sequential effects: superstition or rational behavior? Adv. Neural Inf. Process. Syst. 21, 1873–80 (2009).

31. Cohen, J. D., McClure, S. M. & Yu, A. J. Should I stay or should I go? How the human brain manages the trade-off between exploitation and exploration. Philos. Trans. R. Soc. Lond. B. Biol. Sci. 362, 933–42 (2007).

32. Doya, K. Modulators of decision making. Nat. Neurosci. 11, 410–6 (2008).

33. Yu, A. J. & Dayan, P. Uncertainty, neuromodulation, and attention. Neuron 46, 681–692 (2005).

34. Kolling, N. et al. Value, search, persistence and model updating in anterior cingulate cortex. Nat. Neurosci. 19, 1280–1285 (2016).

35. Holroyd, C. B. & Yeung, N. Motivation of extended behaviors by anterior cingulate cortex. Trends Cogn. Sci. 16, 122–8 (2012).

36. Berridge, C. W. & Waterhouse, B. D. The locus coeruleus-noradrenergic system: modulation of behavioral state and state-dependent cognitive processes. Brain Res. Brain Res. Rev. 42, 33–84 (2003).

37. Aston-Jones, G. & Cohen, J. D. Adaptive gain and the role of the locus coeruleus-norepinephrine system in optimal performance. J. Comp. Neurol. 493, 99–110 (2005).

38. Usher, M., Cohen, J. D., Servan-Schreiber, D., Rajkowski, J. & Aston-Jones, G. The role of locus coeruleus in the regulation of cognitive performance. Science 283, 549–54 (1999).

39. Jepma, M., Te Beek, E. T., Wagenmakers, E.-J., van Gerven, J. M. A. & Nieuwenhuis, S. The role of the noradrenergic system in the exploration-exploitation trade-off: a psychopharmacological study. Front. Hum. Neurosci. 4, 170 (2010).

40. Eldar, E., Cohen, J. D. & Niv, Y. The effects of neural gain on attention and learning. Nat. Neurosci. 16, 1146–53 (2013).

41. Critchley, H. D., Tang, J., Glaser, D., Butterworth, B. & Dolan, R. J. Anterior cingulate activity during error and autonomic response. NeuroImage 27, 885–895 (2005).

42. Ebitz, R. B., Albarran, E. & Moore, T. Exploration disrupts choice-predictive signals and alters dynamics in prefrontal cortex. Neuron 97, 450–61 (2018).

43. Joshi, S., Li, Y., Kalwani, R. M. M. & Gold, J. I. I. Relationships between pupil diameter and neuronal activity in the locus coeruleus, colliculi, and cingulate cortex. Neuron 89, 221–34 (2015).

44. Jepma, M. & Nieuwenhuis, S. Pupil diameter predicts changes in the exploration-exploitation trade-off: evidence for the adaptive gain theory. J. Cogn. Neurosci. 23, 1587–96 (2011).

45. Urai, A. E., Braun, A. & Donner, T. H. Pupil-linked arousal is driven by decision uncertainty and alters serial choice bias. Nat. Commun. 8, 14637 (2017).

46. Dayan, P. & Sejnowski, T. Exploration bonuses and dual control. Mach. Learn. 2, 5–22 (1996).

47. Badre, D., Doll, B. B., Long, N. M. & Frank, M. J. Rostrolateral prefrontal cortex and individual differences in uncertainty-driven exploration. Neuron 73, 595–607 (2012).

48. Green, D. M. & Swets, J. A. Signal detection theory and psychophysics, Wiley, New York, NY (1966).

49. Sanborn, A. N. & Chater, N. Bayesian brains without probabilities. Trends Cogn. Sci. 20, 883–893 (2016).

50. Haefner, R. M., Berkes, P. & Fiser, J. Perceptual decision-making as probabilistic inference by neural sampling. Neuron 90, 649–660 (2016).

51. Gershman, S. J. A unifying probabilistic view of associative learning. PLOS Comput. Biol. 11, e1004567 (2015).

52. Beck, J. M., Ma, W. J., Pitkow, X., Latham, P. E. & Pouget, A. Not noisy, just wrong: the role of suboptimal inference in behavioral variability. Neuron 74, 30–9 (2012).

53. Holroyd, C. B. & Coles, M. G. H. The neural basis of human error processing: reinforcement learning, dopamine, and the error-related negativity. Psychol. Rev. 109, 679–709 (2002).

54. Cavanagh, J. F. & Shackman, A. J. Frontal midline theta reflects anxiety and cognitive control: meta-analytic evidence. J. Physiol. Paris 109, 3–15 (2015).

55. Wessel, J. R., Danielmeier, C., Morton, J. B. & Ullsperger, M. Surprise and error: common neuronal architecture for the processing of errors and novelty. J. Neuropsychiatry Clin. Neurosci. 32, (2012).

56. O’Reilly, J. X. et al. Dissociable effects of surprise and model update in parietal and anterior cingulate cortex. Proc. Natl. Acad. Sci. U. S. A. 110, E3660–9 (2013).

57. Warren, C. M., Hyman, J. M., Seamans, J. K. & Holroyd, C. B. Feedback-related negativity observed in rodent anterior cingulate cortex. J. Physiol. 109, 87–94 (2015).

58. Fouragnan, E., Retzler, C. & Philiastides, M. G. Separate neural representations of prediction error valence and surprise: evidence from an fMRI meta-analysis. Hum. Brain Mapp. 39, 2887–906 (2018).

59. Kennerley, S. W., Dahmubed, A. F., Lara, A. H. & Wallis, J. D. Neurons in the frontal lobe encode the value of multiple decision variables. J. Cogn. Neurosci. 21, 1162–1178 (2009).

60. Boorman, E. D., Rushworth, M. F. & Behrens, T. E. Ventromedial prefrontal and anterior cingulate cortex adopt choice and default reference frames during sequential multi-alternative choice. J. Neurosci. 33, 2242–53 (2013).

61. Kolling, N. et al. Value, search, persistence and model updating in anterior cingulate cortex. Nat. Neurosci. 19, 1280–1285 (2016).

62. Kaplan, R. et al. The neural representation of prospective choice during spatial planning and decisions. PLOS Biol. 15, e1002588 (2017).

63. Hyafil, A., Summerfield, C. & Koechlin, E. Two mechanisms for task switching in the prefrontal cortex. J. Neurosci. 29, 5135–5142 (2009).

64. Amiez, C., Joseph, J. P. P. & Procyk, E. Reward encoding in the monkey anterior cingulate cortex. Cereb. Cortex 16, 1040–55 (2006).

65. Holroyd, C. B. & Coles, M. G. H. Dorsal anterior cingulate cortex integrates reinforcement history to guide voluntary behavior. Cortex 44, 548–559 (2008).

66. Boorman, E. D., Behrens, T. E. J., Woolrich, M. W. & Rushworth, M. F. S. How green is the grass on the other side? Frontopolar cortex and the evidence in favor of alternative courses of action. Neuron 62, 733–43 (2009).

67. Kennerley, S. W., Walton, M. E., Behrens, T. E. J., Buckley, M. J. & Rushworth, M. F. S. Optimal decision making and the anterior cingulate cortex. Nat. Neurosci. 9, 940–7 (2006).

68. Tervo, D. G. R. et al. Behavioral variability through stochastic choice and its gating by anterior cingulate cortex. Cell 159, 21–32 (2014).

69. Meder, D. et al. Simultaneous representation of a spectrum of dynamically changing value estimates during decision making. Nat. Commun. 8, 1942 (2017).

70. Farashahi, S. et al. Metaplasticity as a neural substrate for adaptive learning and choice under uncertainty. Neuron 94, 401–414.e6 (2017).

71. Kolling, N., Behrens, T. E. J., Mars, R. B. & Rushworth, M. F. S. Neural mechanisms of foraging. Science 336, 95–8 (2012).

72. Shenhav, A., Botvinick, M. M. & Cohen, J. D. The expected value of control: an integrative theory of anterior cingulate cortex function. Neuron 79, 217–40 (2013).

73. Browning, M., Behrens, T. E., Jocham, G. & Hospital, J. R. Anxious individuals have difficulty learning the causal statistics of aversive environments. 18, 590–596 (2015).

74. Arnsten, A. F. T. & Goldman-Rakic, P. S. Selective prefrontal cortical projections to the region of the locus coeruleus and raphe nuclei in the rhesus monkey. Brain Res. 306, 9–18 (1984).

75. Rajkowski, J., Lu, W., Zhu, Y., Cohen, J. & Aston-Jones, G. Prominent projections from the anterior cingulate cortex to the locus coeruleus in rhesus monkey. Soc. Neurosci. Abstr. 26, 838–15 (2000).

76. Khamassi, M., Quilodran, R. R., Enel, P., Dominey, P. F. & Procyk, E. Behavioral regulation and the modulation of information coding in the lateral prefrontal and cingulate cortex. Cereb. Cortex 25, 3197–3218 (2014).

77. Warren, C. M. et al. The effect of atomoxetine on random and directed exploration in humans. PLOS One 12, e0176034 (2017).

78. Kane, G. A. et al. Increased locus coeruleus tonic activity causes disengagement from a patch-foraging task. Cogn. Affect. Behav. Neurosci. 17, 1073–1083 (2017).

## References

[1] N. Chopin. A sequential particle filter method for static models. Biometrika, 89(3):539–552, 2002.

[2] N. Chopin, P. E. Jacob, and O. Papaspiliopoulos. Smc2: an efficient algorithm for sequential analysis of state space models. Journal of the Royal Statistical Society: Series B (Statistical Methodology), 75(3):397–426, 2013.

[3] N. D. Daw, J. P. O’doherty, P. Dayan, B. Seymour, and R. J. Dolan. Cortical substrates for exploratory decisions in humans. Nature, 441(7095):876, 2006.

[4] R. Deichmann, J. A. Gottfried, C. Hutton, and R. Turner. Optimized epi for fmri studies of the orbitofrontal cortex. Neuroimage, 19(2):430–441, 2003.

[5] A. Doucet, S. Godsill, and C. Andrieu. On sequential monte carlo sampling methods for bayesian filtering. Statistics and computing, 10(3):197–208, 2000.

[6] M. Jenkinson, C. F. Beckmann, T. E. Behrens, M. W. Woolrich, and S. M. Smith. Fsl. Neuroimage, 62(2):782–790, 2012.

[7] F. Lindsten, T. B. Schön, et al. Backward simulation methods for monte carlo statistical inference. Foundations and Trends^®^ in Machine Learning, 6(1):1–143, 2013.

[8] E. Maris and R. Oostenveld. Nonparametric statistical testing of eeg-and meg-data. Journal of neuroscience methods, 164(1):177–190, 2007.

[9] A. McIntosh. The jackknife estimation method. arXiv preprint arXiv:1606.00497, 2016.

[10] S. Palminteri, M. Khamassi, M. Joffily, and G. Coricelli. Contextual modulation of value signals in reward and punishment learning. Nature communications, 6:8096, 2015.

[11] C. P. Robert. Monte carlo methods. Wiley Online Library, 2004.

[12] J. N. Rouder, P. L. Speckman, D. Sun, R. D. Morey, and G. Iverson. Bayesian t tests for accepting and rejecting the null hypothesis. Psychonomic bulletin & review, 16(2):225–237, 2009.

[13] N. Weiskopf, C. Hutton, O. Josephs, and R. Deichmann. Optimal epi parameters for reduction of susceptibility-induced bold sensitivity losses: a whole-brain analysis at 3 t and 1.5 t. Neuroimage, 33(2):493–504, 2006.

